# Adaptive behavioural strategies to seasonal challenges by a semi-urban feral ungulate

**DOI:** 10.1101/2025.03.12.642745

**Authors:** Debottam Bhattacharjee, Kate J. Flay, Hannah. S. Mumby, Alan G. McElligott

## Abstract

Seasonality, or temporal shifts between wet and dry seasons, profoundly affects the behavioural ecology of animals, particularly large herbivores, in (sub-) tropical climatic conditions. Behavioural strategies are crucial for overcoming challenges associated with seasonality. Group-level strategies to seasonality in the wild have received considerable attention, yet little is known about how large herbivores in human-dominated landscapes respond to seasonal challenges. Moreover, focusing solely on groups can obscure how individual animals of different sexes, ages, and personalities respond. We investigated a semi-urban feral population of a large ungulate, the water buffalo (*Bubalus bubalis*) (*N=71*) during a wet and a dry season. Individual body condition was non-invasively assessed to evaluate whether seasonality impacts physiology. To identify behavioural strategies, we collected data on feeding (grazing and browsing) and space use (core area and home range). Further, using a subset (*n=30*), we investigated the associations between personality and space use in the dry season. Body condition scores were highest during the wet season and then declined during the dry season, indicating an impact of seasonality on physiology. Older individuals were more affected than younger ones. Browsing increased during the dry compared to the wet season. While there was no change in core area use, home ranges expanded in the dry compared to the wet season. Regardless of season, females utilised home ranges larger than males. Females exhibiting higher ‘social tension’ personalities had smaller core areas and home ranges than those with lower social tension. We show that buffalo employ behavioural strategies to overcome challenges associated with seasonality, and these strategies are contingent on intrinsic factors: sex, age, and personality. Our findings offer insights into the ecological principles –habitat use, resource utilisation, and competition– that govern the behavioural ecology of herbivores, and have implications for conservation and welfare in increasingly human-dominated and climate-sensitive landscapes.

## 1. Introduction

Seasonality, or the alternating climatic shifts between wet and dry seasons, drives ecological processes, which in turn influences animal survival (Forrest & Miller-Rushing, 2010; Varpe, 2017; Williams et al., 2017). The wet and dry season of (sub-) tropical climatic conditions exposes animals to periods of high and low resource availability, respectively (Thackeray et al., 2016). More specifically, the dry season’s low resource availability (e.g., reduced grass forage) imposes severe physiological and general survival challenges on herbivores in the wild (Owen-Smith et al., 2005; Woodworth et al., 2018). Adaptive behavioural strategies in animals are imperative to cope with challenges associated with seasonality (Abraham et al., 2022). In comparison to the wet season, during (dry) low resource periods, herbivores reduce general activity (Wittemyer et al., 2007), limit physiologically costly events like reproduction (Ogutu et al., 2014; Owen-Smith & Ogutu, 2013), migrate and expand their home ranges (Aikens et al., 2020; Young et al., 2009), and flexibly use feeding strategies, such as grazing and browsing (Staver & Hempson, 2020). While these behavioural strategies have been extensively studied in the wild, little is known about how large herbivores navigate seasonal challenges in complex human-dominated landscapes.

The rapid expansion of human-dominated landscapes in the Anthropocene is a major biodiversity concern (Lewis & Maslin, 2015; Monsarrat et al., 2019). Immediate and long-term anthropogenic impacts, such as habitat fragmentation and loss due to infrastructure development and climate change, can shape the spatial and social ecology of animals (Gaynor et al., 2024). Large herbivores living in (semi-)urban habitats are of great interest as seasonal challenges can have further implications for human-animal conflict (such as road accidents, crop raiding, and disease transmission) and the maintenance of local biodiversity (such as increased density and overgrazing, and also eco-engineering of habitat) (Lundgren et al., 2024; Carpio et al., 2021; David Walter et al., 2011; Honda et al., 2018; Pascual-Rico et al., 2021; Polfus & Krausman, 2012; Wevers et al., 2020). Typically, (semi-)urban animals, including large herbivores, have smaller home ranges than their wild counterparts (O’Donnell & delBarco-Trillo, 2020), and are often highly sensitive (due to low population size) to seasonal challenges and climate change (Hetem et al., 2014; McCain & King, 2014). However, such responses are not necessarily uniform across individuals within a group, as intrinsic factors like sex, age, personality, and even sterilisation status (if animals are managed) can lead to substantial inter-individual variations (Bonnot et al., 2018; Monestier et al., 2016; Spiegel et al., 2017; Williams et al., 2022). Therefore, it is crucial to examine the adaptive behavioural strategies of (semi-)urban large herbivores with regard to seasonality, emphasising the intrinsic individual-level factors. Such knowledge would help devise appropriate management decisions for successful human-animal coexistence.

The Water buffalo (*Bubalus bubalis*) (hereafter, buffalo) is a globally abundant ungulate species (Minervino et al., 2020). Although a few feral and free-ranging buffalo populations exist worldwide, scientific research is predominantly restricted to livestock. Nonetheless, evidence suggests that free-ranging buffalo are sensitive to seasonality and may exhibit adaptive strategies. For example, in the savannahs and floodplains of the Northern Territory in Australia, the space use patterns of non-native buffalo vary between wet and dry seasons (Campbell et al., 2021; Pike et al., 2024), and they show mixed feeding strategies during low resource periods (Bowman et al., 2010). These feral populations have further implications, as they may positively (as eco-engineers) and negatively (through overgrazing and outcompeting other grazers) impact local biodiversity, contingent on the population size (Lundgren et al., 2024; Mihailou & Massaro, 2021; Petty et al., 2007). However, feral populations of buffalo in human-dominated landscapes are rare and have subsequently received relatively little to no attention.

Lantau Island in Hong Kong accommodates a small population (approximately 115 individuals) of feral and free-ranging buffalo, primarily inhabiting semi-natural and semi-aquatic marshlands around human settlements (Yang et al., 2024). This ‘iconic’ marshland population of buffalo has historical, ecological, and socio-cultural values to the Island and its residents (Yang et al., 2024; So & Dudgeon, 2020), but is also threatened due to increasing developmental activities, such as landfill and habitat fragmentation (WWF-Hong Kong, 2021). Furthermore, semi-aquatic marshlands are subject to intense pressure of terrestrialisation (Koshida & Katayama, 2018), especially during the dry season, leading to deterioration in resource quality and quantity for the buffalo. Thus, understanding their behavioural ecology in Hong Kong’s sub-tropical climatic conditions with alternating wet and dry seasons (Niu & Dudgeon, 2011) is valuable in evaluating the adaptive behavioural strategies of large herbivores to seasonality, particularly in human-dominated landscapes, and their potential biodiversity conservation and welfare implications.

We non-invasively assessed buffalo body condition during a wet and a dry season. We hypothesised that body condition scores would be highest in the wet season and then decline in the dry due to the relatively low availability of nutritional grass forage, indicating the physiological challenges posed by seasonality. Accordingly, we expected buffalo to adjust their feeding behaviour and space use patterns (*sensu* Optimal foraging theory, (Bergman et al., 2001)) as adaptive behavioural strategies. In comparison to the wet season, we predicted that (i) the relative frequency of browsing (to grazing) would increase during the dry season, and (ii) buffalo would expand their home range sizes to mitigate resource competition. We also predicted that females, regardless of season, would utilise larger home ranges than (resident-) males as territoriality would constrain space use patterns of males (Bhattacharjee et al., 2024). Given that buffalo are predominantly marshland dwellers and have site fidelity (Campbell et al., 2021; Dudgeon & Corlett, 2011; So & Dudgeon, 2020), we expected no significant changes in core area use with regard to seasonality. In other words, core area expansion is unexpected due to the geographical confinement of semi-aquatic marshlands to certain locations of the study area (WWF-Hong Kong, 2021). Finally, we predicted that buffalo space use patterns during the dry season would be associated with certain personality traits, particularly ‘social tension’ and ‘general dominance’ (Bhattacharjee et al., 2024). Social tension personality trait in feral buffalo was characterised by approaching and avoiding herd members with the exhibition of self-groom (or self-directed) behaviours, whereas, general dominance was defined by displacing herd members to sit and potentially utilise marshland habitats (Bhattacharjee et al., 2024). As traits similar to social tension and general dominance can influence niche specialisation and habitat exploration (Schirmer et al., 2019), we expected a negative relationship between social tension and space use and a positive relationship between general dominance and space use.

## 2. Methods

### 2.1. Study area and species

We conducted observations on a feral and free-ranging water buffalo population on Lantau Island (22°16’14” N 113°57’10” E), Hong Kong. The buffalo population is predominantly present in the semi-natural marshlands of the southern region of Lantau Island (Yang et al., 2024). We studied the buffalo in and around four geographic locations: Lo Wai Tsuen (Pui O) (22°14’32.7” N 113°58’42.3” E), Lo Uk Tsuen (Pui O) (22°14’29.3” N 113°58’31.1” E), Shap Long Kau Tsuen (22°14’22” N 113°59’27” E), and Shui Hau (22°13’19” N 113°54’48” E) (**Figure S1**). All four locations consist of human habitation, semi-natural freshwater marshlands, backwater rivers, and natural vegetation (such as grasslands, woodlands, and shrublands) common to Lantau Island and, in general, Hong Kong (Dudgeon & Corlett, 2011).

We studied 71 adult buffalo (>60% of the population on Lantau Island; female = 50, resident male = 21; age range = 4 to 22 years, median age = 10 years) between July 2023 and March 2024 (**Table S1**). All males studied were part of herds, and the non-herd living solitary males were not observed. This was primarily due to the reasons that finding and observing solitary males is challenging, and they cover large areas without exhibiting site fidelity (personal observation, D.B., 2023-2024). Thus, we only included the territorial resident males in this study (cf. Bhattacharjee et al., 2024). Individual age details were collected from a local non-government citizen group that has documented the Lantau buffalo population’s demographic information (e.g., birth year with identities) for >20 years. All individuals, except two females (one from Lo Uk Tsuen and the other from Shap Long Kau Tsuen) and six males (three from Lo Wai Tsuen, one from Lo Uk Tsuen, and two from Shap Long Kau Tsuen), were sterilised by the Agriculture, Fisheries and Conservation Department of Hong Kong (AFCD) before the beginning of our study (AFCD, 2013). The sterilisation status of these non-sterilised individuals did not change during the study period. Approximately 65% of the individuals had numbered ear tags (administered by the AFCD). We also developed a photo catalogue based on morphological features (horn shape and structure, relative horn length, and scar marks) to ensure reliable identification of the rest of the individuals. Although the majority of the buffalo (i.e., 88%) are sterilised, social interactions among individuals in our study population resemble those of wild and more natural populations in the forms of complex affiliative patterns, physical aggression, territorial defence and guarding, and dominance-rank relationships (Bhattacharjee et al., 2024). The buffalo population on Lantau Island occasionally (2-4 times a week from December to January) receive supplemental food (like hay and sweet potato leaves) from local non-government citizen groups. However, the amount of provisioned food and their nutritional values were unknown.

### 2.2. Ethical note

We obtained ethical approval for our study from the Animal Research Ethics Sub-Committee of City University of Hong Kong (Reference no. AN-STA-00000195). Behavioural observations of buffalo were conducted from a distance of at least 20 m without direct human intervention, and we also adhered to the ethical guidelines of the ASAB/ABS while conducting this study (ASAB Ethical Committee & ABS Animal Care Committee, 2022).

### 2.3. Data collection

We used a transect sampling method to collect data between July 2023 and March 2024. During the transects, we followed predetermined routes at each of the four study locations (Lo Wai Tsuen, Lo Uk Tsuen, Shap Long Kau Tsuen, and Shui Hau). Upon sighting an individual buffalo, we recorded the following: (i) body condition (Ezenwa et al., 2009), by visually assessing and scoring four regions of the body (ribs, tail, hips, and spine), along with coat condition (see below), (ii) feeding behaviour, whether an individual was feeding by grazing on grass or browsing on shrubs or trees or not feeding, and (iii) location, using a mobile GPS device on *Google My Maps* software’s ‘satellite’ basemap.

Data on body condition and feeding behaviour were recorded using a handheld video camera (Panasonic HC-V785), while GPS locations of individuals were collected live. The duration of the recording depended on the area size and the number of individuals sampled (range of cumulative duration = 3 min to 14 min). We conducted only one transect per study location on a given day, but multiple study locations were often sampled on the same day. Additionally, we did not conduct transects within a given study location on consecutive days to avoid potential data dependence, as buffalo are usually slow-moving and exhibit site fidelity (Campbell et al., 2021). The sampling time was pseudo-randomised for each study location, ensuring the data represented a broad time window between 0930 and 1730 hours. We conducted 1 - 3 transects per week over the entire study period. During a transect, the same roads in all study locations were walked more than once to cover the entire area (Bhattacharjee et al., 2024). However, data on an already sampled individual was not recorded twice. We set a criterion of collecting 80 data points per individual over the entire study (Börger et al., 2006; Laver & Kelly, 2008), specifically for feeding behaviour and individual GPS locations. Therefore, if an individual was not sighted during a transect, we conducted additional ones on the following days to fulfil the criterion exclusively for the given individual. Individuals who had already been sampled during the first transects were not considered during those additional transects. We calculated the percentages of finding individuals during the first transects (range = 87.5% - 100%; see **Table S2**). Body condition was assessed monthly, i.e., nine times during the study period between July 2023 and March 2024.

### 2.4. Characterising wet and dry seasons

We extracted monthly mean air temperature (in °C) and total rainfall (in mm) data between July 2023 and March 2024 from the Hong Kong Observatory public database (Hong Kong Observatory, 2024). To determine the wet and dry months, we conducted a *k-means clustering* analysis based on temperature and rainfall data. We used a *k value* of 2, i.e., considering potential wet and dry seasons. The two clusters (cluster one: July to October; cluster two: November to March) explained >80% of the variance of the data (**Figure S2**). Cluster one was characterised by median temperature and rainfall of 29.4°C (range: 26.4°C – 30.1°C) and 391.2 mm (range: 175.2 mm – 546 mm), respectively. Cluster two, by contrast, was characterised by median temperature and rainfall of 19.4°C (range: 17.9°C – 23.5°C) and 4.1 mm (range: 0.9 mm – 21.6 mm), respectively. We further collected the temperature and rainfall data between July 2019 and March 2023 to check whether our study years resembled those of previous years’ weather patterns. We found consistent patterns, i.e., higher temperatures and rainfall between July and October than between November and March (Mann Whitney U test – Temperature: *Z* = -5.07, *p* <0.001; Rainfall: *Z* = -4.66, *p* <0.001). Subsequently, we determined July to October (or cluster one) as the wet season and November to March (or cluster two) as the dry season. Body condition, feeding behaviour, and space use data were divided accordingly into wet and dry seasons for statistical analyses.

### 2.5. Data coding, preparation, and analysis

#### 2.5.1. Body condition scoring

Body condition scoring is a widely accepted and validated non-invasive indicator for welfare and fitness-related traits in herbivores (Parker et al., 2009; Powell et al., 2024). A standardised non-invasive body condition scoring technique, originally used for African buffalo (*Syncerus caffer caffer*), was employed in our study (Ezenwa et al., 2009). Although a comprehensive scoring technique has been developed for and implemented in water buffalo (Alapati et al., 2010), the free-ranging living conditions did not allow us to inspect the individuals from a close distance (i.e., <20 m) necessary for the method. In our currently implemented method, four regions of the body (ribs, tail, hips, and spine; **Figure S3**), along with coat condition, were scored separately on a scale of 1 to 5, with higher scores indicating a better condition (cf. (Ezenwa et al., 2009)). Based on all five measures, a composite body condition score (i.e., the overall body condition) was calculated with a maximum potential value of 25. For a general understanding, we describe the two extreme scores for each region of the body considered for scoring: (i) ribs – a score of 1 indicates clearly visible ribs with deep depression between them, while a score of 5 indicates invisible ribs due to the presence of fatty layers, (ii) tail – a score of 1 indicates the tissue surrounding tail base forms round hollows, whereas 5 indicates that the tail base sits in a depression surrounded by fatty tissues, (iii) hips – a score of 1 suggests that the hip bones protrude beyond hip points, while a score of 5 indicates no visually apparent hip bones, (iv) spine – a score of 1 indicates visually distinguishable vertebrae, whereas 5 indicates invisible vertebrae due to the presence of fatty layers, and finally (v) coat – a score of 1 indicates bald or sparsely coated body, whereas 5 suggests the presence of glossy coat with no bald patches. In the original study by Ezenwa et al. (2009) on African buffalo (*Syncerus caffer caffer*), coat condition was dropped from the composite body condition measure as it was not sufficiently validated. However, we decided to retain it in our calculation of overall body condition score as the wet (average ± standard deviation = 4.93 ± 0.25) and dry (4.76 ± 0.42) season coat conditions were statistically different (Mann Whitney U test: *p* <0.001). This suggests that coat condition likely had an impact on the overall composite measure of body condition in feral water buffalo in Hong Kong.

Due to the subjective nature of the body condition scoring technique, we performed a reliability test between the main observer (D.B.) and another trained person who was unaware (i.e., blind coder) of the study objectives. Based on 16% of the body condition data, we obtained a *Cohen’s kappa* value of 0.94, indicating a high reliability. We pooled the monthly body condition scores and calculated averages for the wet (based on four data points per individual) and dry (based on five data points per individual) seasons.

#### 2.5.2. Feeding behaviour

We coded the occurrences of feeding from the videos, focusing on two strategies: grazing and browsing. Grazing was recorded when an individual was observed to place their mouth at the level of grass with clear visible contact, along with visible movement of the mouth (**Figure S4a**). Browsing was defined as an individual standing with their mouth visibly in touch with plant leaves or shrubs but not grass, along with the visible movement of the mouth (**Figure S4b**). Notably, during a transect, only the first instance was coded to determine whether an individual was feeding (either by grazing or browsing) or not. We pooled and counted all occurrences of individual-level feeding behaviour during the transect sampling and then calculated the proportion of browsing.

#### 2.5.3. Space use

Individual geo-locations were used to investigate the space use patterns of buffalo during the wet and dry seasons. We calculated home ranges and core areas for individuals by computing minimum convex polygon (MCP) and Kernel density estimation (KDE). As MCP and KDE often over- or underrepresent space use patterns contingent on the number and distribution of data points (i.e., locations), it is recommended to use more than one method to check validity; we computed 100% MCP, 95% KDE with both reference (HREF) and least squares cross-validation (LSCV) bandwidth selectors, and 50% KDE with both HREF and LCSV bandwidth selectors (Börger et al., 2006; Laver & Kelly, 2008; Lichti & Swihart, 2011). A minimum sample size of 30 is recommended for the LSCV method (Seaman et al., 1999), whereas the HREF method can be applied based on at least a sample size of 10 (Webb et al., 2011). We used 40 individual geo-locations each for wet and dry seasons in the current study. For MCP, a convex hull was computed around the spatial points, showing the spatial extent of the data. The computed convex hull was plotted to check the spatial boundary of the dataset visually. KDE was computed to estimate the utilisation distribution, with two specific contours extracted: 95% and 50%, representing the broader home range and core area, respectively. The resulting contours were overlaid on a plot of the spatial points to visualise the distribution of the observed locations and the estimated home ranges and core areas. Home range and core area values were calculated in square metres and converted into square kilometres.

#### 2.5.4. Personality

To investigate whether personality traits predict the space use patterns in buffalo, we utilised a published dataset on personality, representing a subset (30 adult females) of our study population (Bhattacharjee et al., 2024). Based on focal observations spanning a wet and a dry season, followed by a ‘bottom-up’ approach, the study reported three consistent inter-individual traits in female buffalo: *social tension* (positively correlating behaviours: approaching a conspecific, avoiding a conspecific, and self-groom), *vigilance* (vigilance behaviour towards a within-herd event or herd member), and *general dominance* (inversely associated behaviours of sitting and displacing a herd member). We used individual scores for each personality trait and separately investigated their associations with the KDE (i.e., 95% and 50% KDE with LSCV bandwidth selector) values for the dry season.

### 2.6. Statistics

All statistical analyses were performed using R (version 4.4.1) (R Development Core Team, 2019). MCP was calculated using the *sf* and *sp* packages (Pebesma, 2018; Roger et al., 2013), and KDE was calculated using the *adehabitatHR* package (Calenge, 2006). We applied a Bayesian mixed modelling approach for analyses using the *brms* package (Bürkner, 2018). We used weakly informative priors to guide the estimation process while allowing the data to dominate the posterior distributions (Lemoine, 2019; McElreath, 2020). We used the *correlation* package to conduct partial Bayesian correlations (Makowski et al., 2020).

We built five models (*Models 1 – 5*) to investigate the following: buffalo body condition (*Model 1*), proportion of browsing (i.e., occurrences of browsing divided by total occurrences of feeding through grazing and browsing) (*Model 2*), home range size using MCP (*Model 3*), home range size using 95% KDE with LSCV bandwidth selector (*Model 4*), and core area size using 50% KDE with LSCV bandwidth selector (*Model 5*). Notably, KDE with HREF bandwidth selectors consistently, i.e., for both 95% and 50% estimates, visually overestimated home range and core area in comparison to MCP and KDE with LSCV bandwidth selectors (**Figure S5**). Additionally, since KDE with LSCV estimate is a more highly recommended method for assessing animal range use than KDE with HREF, we avoided using the latter in our analyses (Powell, 2000). In all models, we included season (wet and dry), age, and sex (female and male) as predictors, and individual identities nested within the herds as random effects.

In Model 1, a Gaussian distribution was used, and a normal prior with a mean (m) of 0 and a standard deviation (SD) of 2 was included for the intercept term. We applied a normal prior for all other fixed-effect coefficients with m = 0 and SD = 1. In Model 2, we used a binomial distribution and identical normal priors for the intercept and fixed effects to that of Model 1. The Gaussian Model 3 included a normal prior with m = 0 and SD = 5 for the intercept term. The normal prior for all other fixed effects was identical to Models 1 and 2. We log-transformed the MCP data to reduce its positive skewness. The priors, m, and SD in Model 4 were identical to Models 1 and 2. Log transformation of the 95% KDE (with LSCV) estimates was performed here. Model 5 was identical to Model 4, except a square root transformation was used for the 50% KDE (with LSCV) data. In all models (Models 1 – 5), an exponential prior with a rate parameter of 1 was assigned to the SD of the random effects. For better convergence, models were subjected to 5500 – 7000 iterations with 1000 – 1500 warmup sessions. Finally, we performed partial Bayesian correlations with the Spearman method to investigate the associations between the three personality traits (social tension, vigilance, and general dominance) and KDE (95% and 50% with LSCV) exclusively for the dry season.

We reported median estimate coefficients (*Est*), 89% credible interval (*crl*) that contains 89% of the posterior probability density function, and probability of direction (*pd*), indicating certainty of an effect. We established model convergence by following Bayesian statistics guidelines (Depaoli & van de Schoot, 2017). For all models, we checked trace and autocorrelation plots, Gelman-Rubin convergence estimations, and density histograms of posterior distributions. We sampled posterior distributions using 10,000 iterations and 2000 warmups. Furthermore, we used the “*DHARMa.helpers*” package (Rodríguez-Sánchez, 2024) and investigated model fits by examining the residual distributions (i.e., deviation, dispersion, and outliers).

## 3. Results

### 3.1. Body condition score

Buffalo body condition scores were highest during the wet season (mean ± standard deviation = 23.87 ± 0.88) and then declined in the dry season (19.20 ± 1.30), suggesting a strong effect of season on physiology (*Est* = -1.58, 89% *crl* = [-1.68, -1.49], *pd* = 1.00; **Figure 1, Figure S6**). Age also predicted individual body condition (*Est* = -0.10, 89% *crl* = [-0.12, -0.07], *pd* = 1.00; **Figure 1**). The negative relationship indicates that body condition scores declined with increasing age of the individuals. No effect of sex was found (*Est* = - 0.01, 89% *crl* = [-0.14, 0.12], *pd* = 0.56; **Figure 1**).

**Figure 1.**
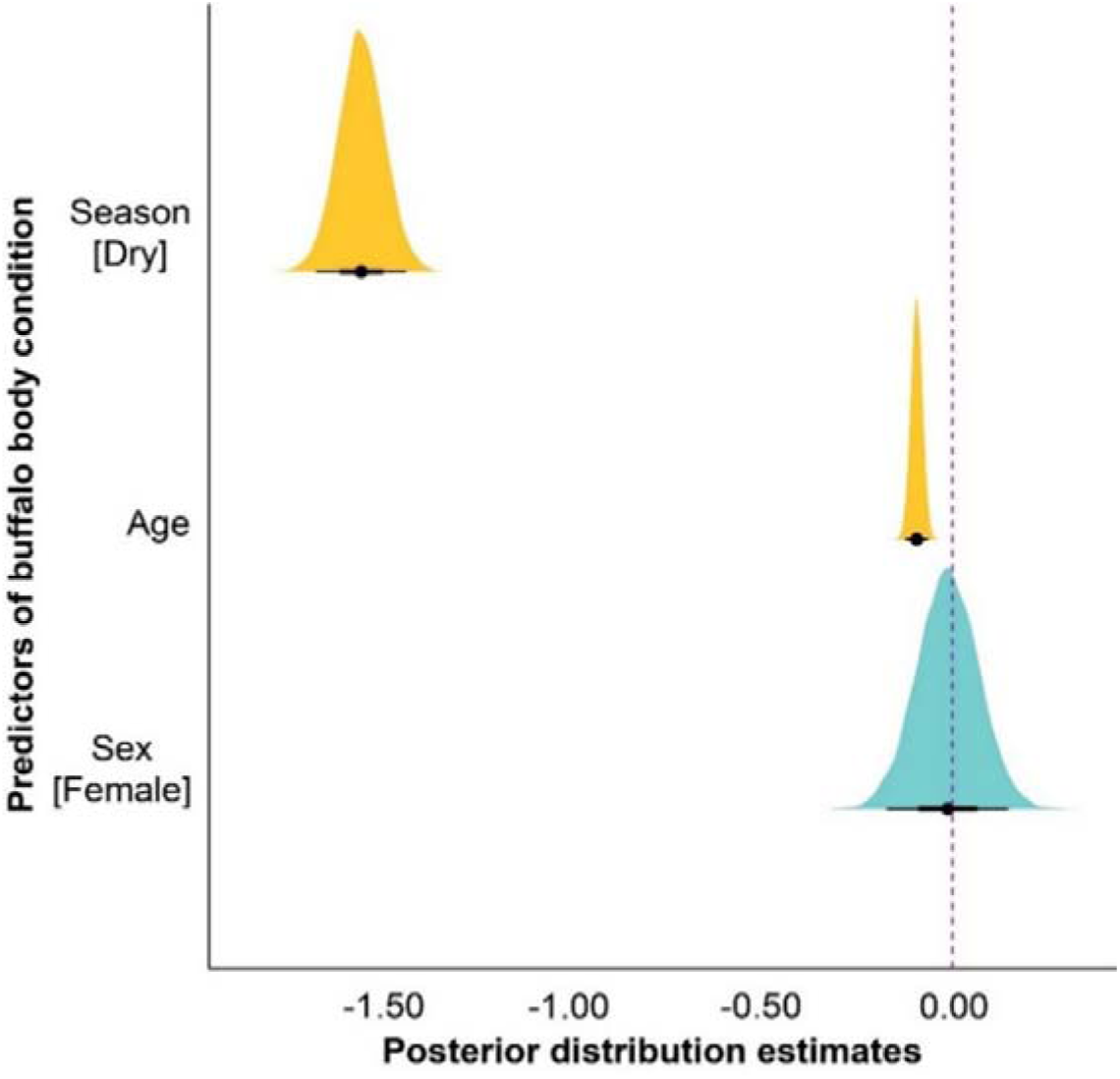
Effects of season, age, and sex on buffalo body condition. The posterior predictive effects of season, age, and sex on buffalo body condition scores (*Model 1*). Yellow colours indicate robust effects of the corresponding predictors, while the blue colour indicates no effect. The width of the ‘half-eye’ represents data distribution (89% *crl*), and solid black points on the horizontal bars indicate the median values. The vertical dashed line indicates a parameter estimate of zero. An overlap of the *crl* values of the predictors with this line suggests no effect.

### 3.2. Feeding behaviour

Grazing was the predominant feeding strategy during the wet season (percentage feeding: grazing = 99.95%, browsing = 0.05%), whereas a mixed strategy (grazing = 96.82%, browsing = 3.18%) was observed in the dry season. The proportion of browsing frequency increased during the dry season (0.83 ± 1.24) compared to the wet (0.01 ± 0.11) season (*Est* = 1.69, 89% *crl* = [1.22, 2.25], *pd* = 1.00; **Figure 2**). There was no effect of age (*Est* = 0.04, 89% *crl* = [-0.02, 0.10], *pd* = 0.86) or sex (*Est* = -0.00, 89% *crl* = [-0.34, 0.33], *pd* = 0.50) on feeding behaviour (**Figure 2**).

**Figure 2.**
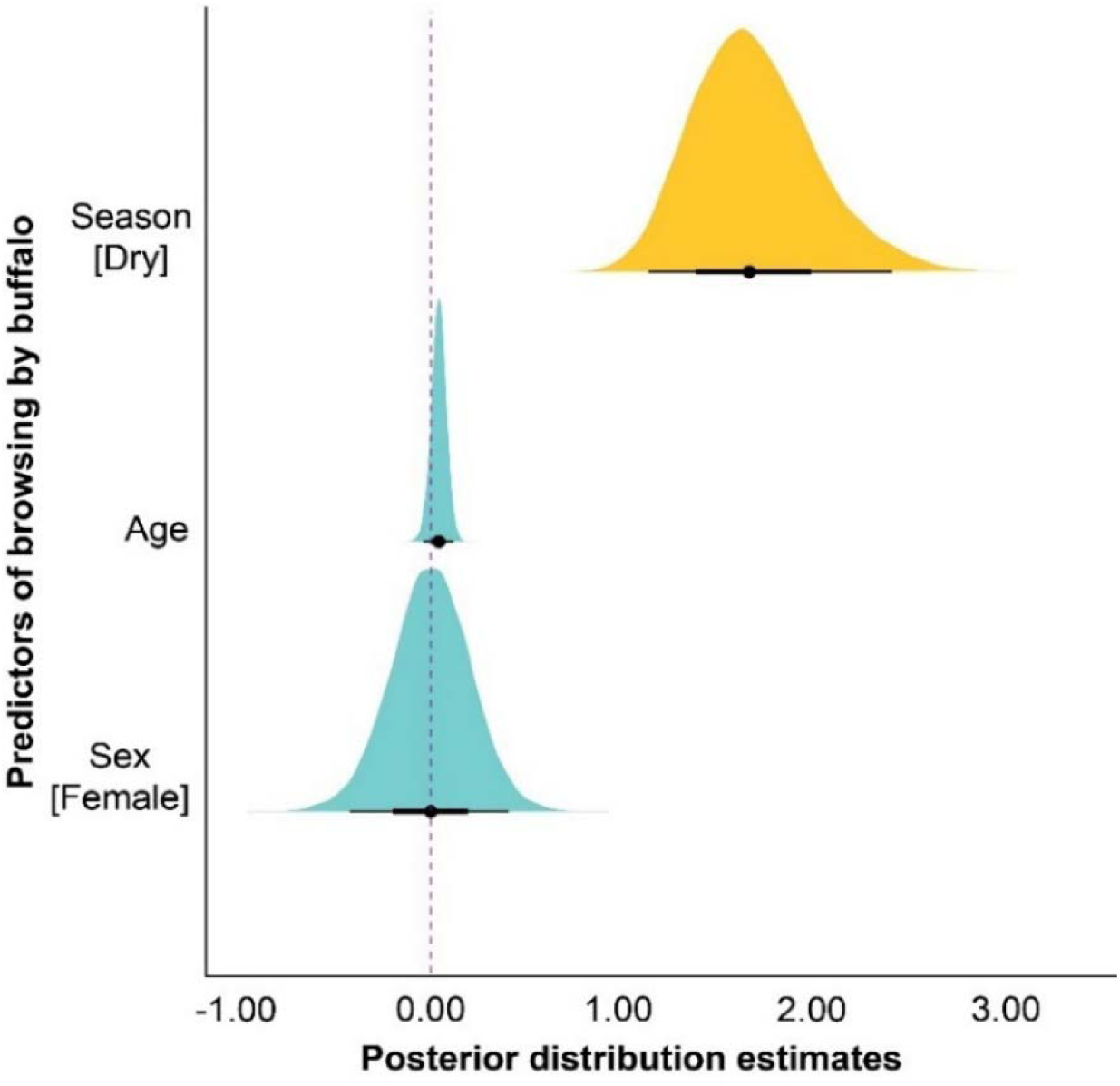
Effects of season, age, and sex on feeding behaviour of buffalo. The posterior predictive effects of season, age, and sex on the proportion of browsing (to grazing) (*Model 2*). Yellow colour indicates robust effects of the corresponding predictor, while blue colours indicate no effect. The width of the ‘half-eye’ represents data distribution (89% *crl*), and solid black points on the horizontal bars indicate the median values. The vertical dashed line indicates a parameter estimate of zero. An overlap of the *crl* values of the predictors with this line suggests no effect.

### 3.3. Space use patterns

We found variations in space use patterns between wet and dry seasons. See **Figure S7** for the location-specific distribution overview of buffalo during wet and dry seasons.

#### 3.3.1. Home range using MCP

The home range of buffalo was 0.09 ± 0.05 km^2^ during the wet season, which increased more than twice to 0.19 ± 0.20 km^2^ in the dry season, suggesting a strong effect of season (*Est* = 0.13, 89% *crl* = [0.11, 0.16], *pd* = 1.00; **Figure 3a**). We did not find any effect of age on home range (*Est* = -0.00, 89% *crl* = [-0.01, 0.00], *pd* = 0.82; **Figure 3a**). However, a strong effect of sex was found (*Est* = 0.23, 89% *crl* = [0.20, 0.27], *pd* = 1.00; **Figure 3a**). Females (0.18 ± 0.17 km^2^) utilised close to four times larger home range than males (0.04 ± 0.03 km^2^), irrespective of season.

**Figure 3.**
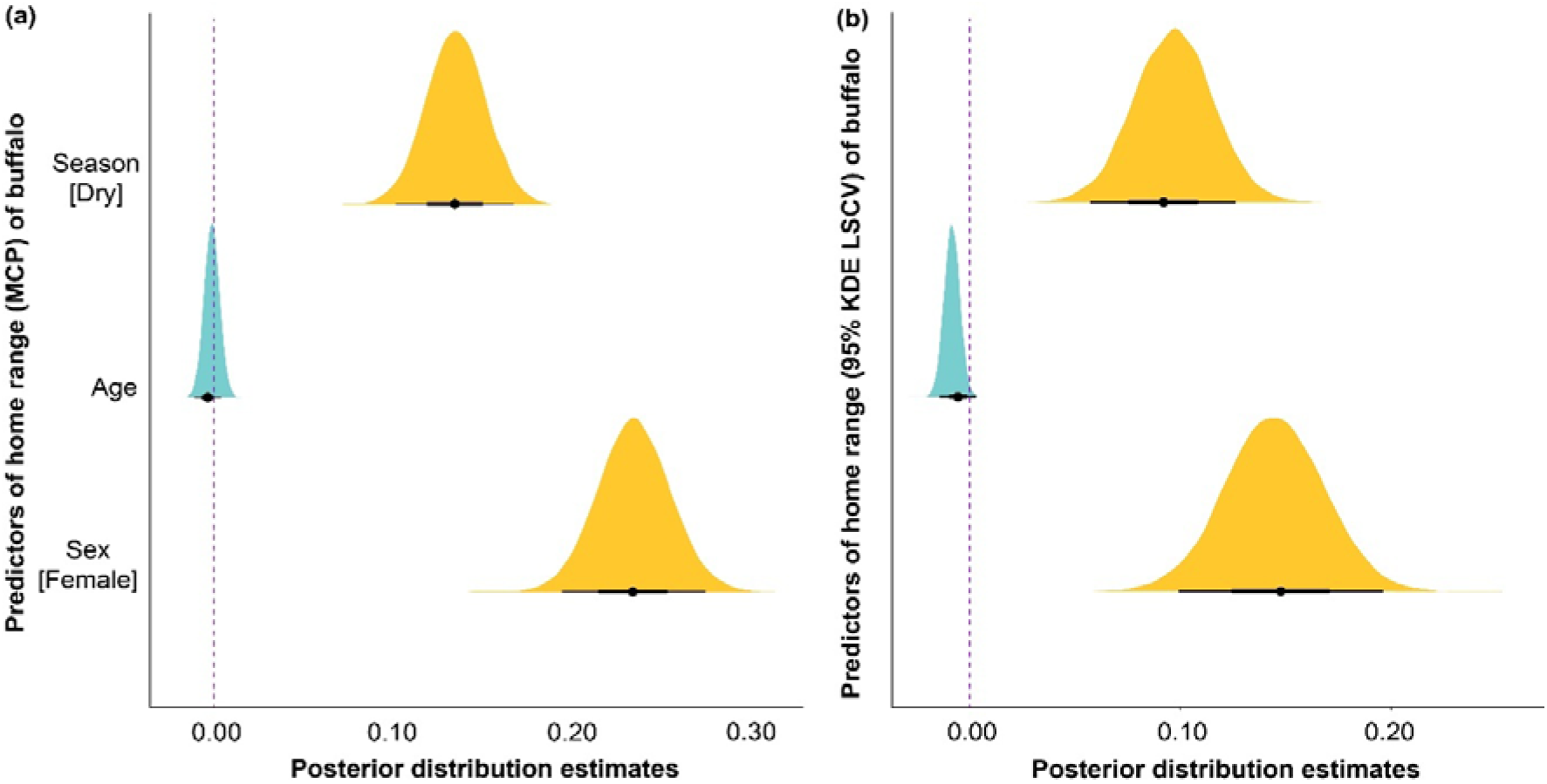
Effects of season, age, and sex on space use by buffalo. **(a)** The posterior predictive effects of season, age, and sex on buffalo home range using MCP method (*Model 3*). (b) The posterior predictive effects of season, age, and sex on buffalo home range using 95% KDE method with LSCV bandwidth selector (*Model 4*). Yellow colours indicate robust effects of the corresponding predictors, while the blue colour indicates no effect. The width of the ‘half-eye’ represents data distribution (89% *crl*), and solid black points on the horizontal bars indicate the median values. The vertical dashed line indicates a parameter estimate of zero. An overlap of the *crl* values of the predictors with this line suggests no effect.

#### 3.3.2. Home range using 95% KDE LSCV

Since MCP and 95% KDE with LCSV bandwidth selector estimated buffalo home ranges similarly (and unlike 95% KDE with HREF, cf. **Figure S5**), we investigated whether the seasonal influence on home range pattern was consistent (cf. Model 4). Home range size was 0.11 ± 0.09 km^2^ in the wet and 0.15 ± 0.10 km^2^ in the dry season. Thus, similar to MCP, we found a strong effect of season on buffalo home range calculated using 95% KDE with LSCV bandwidth selector (*Est* = 0.09, 89% *crl* = [0.06, 0.12], *pd* = 1.00; **Figure 3b**). No effect of age was found (*Est* = -0.01, 89% *crl* = [-0.01, 0.00], *pd* = 0.88; **Figure 3b**). Like MCP, a strong effect of sex was found here (*Est* = 0.15, 89% *crl* = [0.11, 0.19], *pd* = 1.00; **Figure 3b**). Females (0.17 ± 0.10 km^2^) utilised more than three times bigger home range than males (0.05 ± 0.03 km^2^), irrespective of dry and wet seasons.

#### 3.3.3 Core area using 50% KDE LSCV

Core areas were 0.03 ± 0.02 km^2^ in dry and 0.026 ± 0.02 km^2^ in wet seasons. Regardless of season, areas were 0.03 ± 0.02 km^2^ for females and 0.012 ± 0.007 km^2^ for males. We did not find any evidence of season (*Est* = 0.47, 89% *crl* = [-1.08, 2.01], *pd* = 0.68), age (*Est* = 0.41, 89% *crl* = [-0.64, 1.48], *pd* = 0.73), or sex (*Est* = 1.17, 89% *crl* = [-0.42, 2.75], *pd* = 0.88) influencing or being associated with the core area use.

As sterilisation status can influence core area use, we compared the sterilised and non-sterilised males in an additional model (identical priors to *Model 5*). Due to the relatively small number of non-sterilised males (i.e., 6 out of 21 males), we pooled the season data. Non-sterilised males showed a general tendency (*Est* = 0.01, 89% *crl* = [0.00, 0.03], *pd* = 0.98) of using larger core areas (0.017 ± 0.009 km^2^) than sterilised (0.009 ± 0.005 km^2^) males.

### 3.4 Association of space use in dry season with personality

We found a moderately strong negative correlation between social tension and 95% KDE home range (*rho* = -0.39, n = 30, 89% *crl* = [-0.62, -0.10], *pd* = 0.99), suggesting that female buffalo with higher social tension had smaller home range than those with lower social tension. No associations were found for the vigilance (*rho* = 0.002, n = 30, 89% *crl* = [-0.30, 0.31], *pd* = 0.51) and general dominance traits (*rho* = -0.28, n = 30, 89% *crl* = [-0.54, 0.02], *pd* = 0.91). Similarly, we found a strong negative correlation between social tension and 50% KDE core area estimate (*rho* = -0.40, n = 30, 89% *crl* = [-0.63, -0.11], *pd* = 0.98), suggesting that female buffalo with higher social tension had smaller core area than those with lower social tension. Consistent with the home range results, no associations were found for the vigilance (*rho* = 0.0002, n = 30, 89% *crl* = [-0.31, 0.31], *pd* = 0.53) and general dominance traits (*rho* = -0.24, n = 30, 89% *crl* = [-0.51, 0.07], *pd* = 0.86).

## 4. Discussion

Resources that fluctuate seasonally bring survival challenges to herbivores, which may increase manifold when animals live in fragmented human-dominated landscapes (Dagtekin et al., 2024). Little is known about how large herbivores adaptively respond to seasonal challenges in such landscapes. We investigated a population of semi-urban feral water buffalo living under sub-tropical climatic conditions with alternating wet and dry seasons. Our results demonstrate that buffalo living in this human-dominated landscape were highly sensitive to seasonality and faced physiological challenges, particularly during the low resource period or dry season. As predicted, we observed a decline in body condition between the wet and the dry season. Adaptive behavioural strategies, in the form of increased browsing and expanded home ranges, were found in the dry season compared to the wet. In line with our predictions, intrinsic factors such as sex, age, and, to an extent, personality were associated with the physiological impact of seasonality and subsequent behavioural responses. While we discuss these findings in light of the broader ecological principles of habitat use, resource utilisation, and competition at the spatial-social interface, we also emphasise the biodiversity conservation and welfare implications of our findings (Webber et al., 2023).

### 4.1 Seasonality and body condition scoring

Both intrinsic (e.g., sex, age, and disease) and extrinsic (e.g., environmental seasonality) factors can influence physiology and, subsequently, an individual’s body condition (Rödel et al., 2023; Stephenson et al., 2020). We found independent effects of age and seasonality on buffalo body condition scores. While age is a strong predictor of general health, with older individuals less able to cope due to senescence (Rödel et al., 2023), a lower average score during the dry season (in comparison to the wet season) highlights the adverse effects of seasonality, thus potentially heightened physiological stress (Pokharel et al., 2017). Nevertheless, the relatively high average body condition score, which was sustained during the wet season, indicates that our study population is unlikely to be under density-dependent resource competition pressure (Gaidet & Gaillard, 2008) and has access to a sufficiently suitable quality habitat (Ranglack & du Toit, 2015). Furthermore, a visually sharp decline in body condition was observed after the first two months of the dry season (cf. **Figure S6**). This pattern could be attributed to the residual effects of rainfall in the late wet season on grass growth and potentially low density-dependent competition.

### 4.2 Seasonality and feeding behaviour

A mixed feeding strategy, i.e., a flexible switch from grazing to browsing, particularly during resource-limited periods (often called seasonal dietary shifts), can be beneficial (Staver & Hempson, 2020) and has been reported in several herbivore species (Abraham et al., 2019; Codron et al., 2007). While not essentially a mixed feeding strategy, in comparison to the wet season, we found an >80-fold increase in browsing during the dry season (3.13% increase based on overall feeding budget) in our observed population of buffalo. The null effects of sex and age indicate that this seasonal shift was a strategy shared across the entire population. African (*Syncerus caffer*) and Asian buffalo are often considered generalist bulk grazers; however, several studies have highlighted the variable nature of grazing with a preference for forage quality over quantity (Kaszta et al., 2016) and regional and seasonal dietary shift strategies (Beekman & Prins, 1989; Bowman et al., 2010; Codron et al., 2007).

In contrast to the dry season, we found only one instance of browsing during the wet season, thus indicating an exclusive seasonal dietary shift in the feral buffalo of Hong Kong. This does not translate into more frequent browsing than grazing, as the latter was still the predominant feeding strategy (browsing = 3.18% and grazing = 96.82%) in the dry season. A more nuanced and informative dietary compositional pattern in feral Hong Kong buffalo could be achieved by investigating faecal samples and applying stable carbon isotopes (i.e., C_3_ for browse and C_4_ for graze; Codron et al., 2007) or DNA metabarcoding techniques (Kartzinel et al., 2015). Additionally, an assessment of the nutritional values of forage with regard to body mass would be an interesting future study that would allow us to better understand the influence of seasonality on feeding behaviour and general welfare conditions of buffalo. Finally, a predominantly high grazing strategy can have implications for local biodiversity. Bulk grazing by buffalo can facilitate the growth of less abundant plant species (Lundgren et al., 2024). Therefore, an assessment of plant diversity between buffalo grazing and non-grazing zones at a restricted spatial scale would be valuable in determining their impact on biodiversity.

### 4.3 Seasonality and space use

Adjusting space use patterns by large herbivores in response to seasonally fluctuating resources is arguably the most prominent adaptive behavioural strategy. Large herbivores, like wildebeest (*Connochaetes taurinus*), migrate in search of food during harsh environmental conditions (Holdo et al., 2009); however, even non-migratory species with high site fidelity exhibit changes (e.g., by expansion of home range) in how they utilise their habitat during the dry season (Bukombe et al., 2018). In a spatially constrained human-dominated landscape on Lantau Island (WWF-Hong Kong, 2021), we found seasonal adjustments in space use by buffalo, such as an increase in the area of their home range during dry compared to the wet season. This pattern of change in home rage is consistent with wild populations of bovids, such as the African buffalo which cover a higher average daily distance in the dry season than in the wet (Roug et al., 2020). Similarly, Asian water buffalo in the floodplains of Northern Australia increase movements and space use during the dry season (Campbell et al., 2021). In contrast, a recent study reported larger home ranges in Asian water buffalo during the wet season in comparison to the dry season (Pike et al., 2024). This contrasting pattern can be attributed to the contraction of wetlands in the habitat, which led individuals to optimise their resources during the dry season within a restricted space. Additionally, buffalo living in the savannahs of Northern Australia have comparable space use patterns between wet and dry seasons (Campbell et al., 2021). These findings clearly suggest that the variations in available resources and habitat type drive space use patterns. Yet, compared to these studies on the same or related species of buffalo, we found the home range of individuals of our feral buffalo population to be much smaller. Such a pattern is to be expected in species present in human-dominated landscapes (O’Donnell & delBarco-Trillo, 2020), where habitats are highly fragmented due to anthropogenic activities.

As predicted, space use patterns in our study buffalo population were associated with intrinsic individual-level factors. Resident males utilised smaller home ranges than females, irrespective of seasons. This difference could potentially be due to a constraint of male territoriality (Stammes et al., *unpublished data*). Expanding their home range would require a resident male to cross other resident males’ territories, which could lead to costly physical encounters. Adult males generally have higher social status than females within a herd, as evidenced by frequent guarding of females and unidirectional aggressive instances towards females (personal observation, D.B., 2023-2024, also see (Bhattacharjee et al., 2024)). Therefore, males may benefit from reduced intra-specific resource competition during the dry season when females expand their home ranges. While the core areas were found to be comparable across seasons, there was a tendency for non-sterilised males to utilise a larger core area than sterilised males. Although the effect size was not large due to the limited number of non-sterilised males in our sample (6 of 21), this could be investigated in the future by including more non-sterilised males. The temporally stable close social associations (Bhattacharjee et al., 2024) among females may prevent them from escaping competition, where home range expansion during the dry season is necessary. Nonetheless, we found no overlap of expanded home ranges among individuals from the four herds, even for the relatively adjacent ones, with no physical barriers preventing them from mixing.

We found partial support for our hypothesis of personality traits influencing space use patterns. In our study, general dominance did not appear to be associated with space use patterns in females, but social tension and space use patterns were negatively associated. We suspect that relatively higher social tension in females could indicate a reluctance to be bold and explorative, leading to reduced space utilisation (Stiegler et al., 2022). In line with this, it would also be important in the future to investigate the association between male personalities and space use patterns, focusing primarily on territoriality.

#### 4.3.1 Seasonal space use and biodiversity implications

Seasonal space use patterns by buffalo have implications for local biodiversity. Feral buffalo on Lantau Island live in and around semi-natural marshlands (Yang et al., 2024; So & Dudgeon, 2020). Semi-natural marshlands are considered a highly productive and alternate ecosystem to natural wetlands as they accommodate a wide range of aquatic macroinvertebrates (Natuhara, 2013). However, these marshlands are susceptible to terrestrialisation, a transformation process through which the diversity of aquatic macroinvertebrates can perish (Koshida & Katayama, 2018). Buffalo, and in general bovids, through grazing, trampling, and wallowing (exclusive to buffalo), can halt this process of terrestrialisation and maintain a high biodiversity, and even help restore ecosystems (So & Dudgeon, 2020; Bora et al. 2024). Note that the core area of buffalo fully overlapped with the distribution of marshlands on Lantau Island ((WWF-Hong Kong, 2021); also see **Figure S7**), which also remained comparable between wet and dry seasons. Thus, the marshlands can be considered high-activity zones of buffalo on Lantau Island. Experimental studies in future would be useful to establish the role of buffalo on marshland biodiversity more firmly (see Perrino et al., 2021; Georgoudis et al., 1999). Nevertheless, protecting the severely threatened marshlands using buffalo as agents should be an immediate first step (cf. (So & Dudgeon, 2020; WWF-Hong Kong, 2021)). This might also reduce potential buffalo-human conflicts on the island (Tang, 2017), especially during the dry season range expansion.

## 5. Conclusion

In conclusion, we provide compelling evidence based on observations that buffalo, a large herbivore species, living in a human-dominated landscape, exhibit sensitivity to seasonality. In particular, the dry season poses physiological challenges, leading to altered feeding behaviour and space use patterns by buffalo. We also show that these adaptive behavioural strategies vary with sex, age, and personality, and thus, seasonality may not affect a population uniformly. However, our study has two limitations. First, tracking individuals using only direct observations and not employing more precise automated tools like GPS collars limits our data collection to areas where animals are visible. This may bias our sampling by underrepresenting buffalo use of dense habitats such as woodlands. However, note that a very high proportion of individuals were found during the first transects (cf. **Table S2**). Second, we were unable to quantify the impacts of supplemental feeding on natural feeding and space use patterns of buffalo. While we acknowledge that such an influence is inevitable, the robust evidence of alternative strategies (i.e., both feeding and space use) suggests that the natural behavioural tendencies of buffalo were not suppressed. Nonetheless, future studies should tackle the two identified shortcomings, i.e., by using precise tracking tools and accounting for supplement feeding impacts, to add more to our already valuable findings regarding the behavioural ecology of buffalo in human-dominated landscapes.

## Author contributions

Conceptualisation: Debottam Bhattacharjee, Kate J. Flay, Hannah S. Mumby, and Alan G. McElligott; Data curation: Debottam Bhattacharjee; Methodology: Debottam Bhattacharjee, Kate J. Flay, Hannah S. Mumby, and Alan G. McElligott; Software and visualisation: Debottam Bhattacharjee; Funding acquisition: Alan G. McElligott, Kate J. Flay, and Hannah S. Mumby; Project administration: Alan G. McElligott; Writing—original draft: Debottam Bhattacharjee; Writing—review and editing: Debottam Bhattacharjee, Kate J. Flay, Hannah S. Mumby, and Alan G. McElligott. All authors contributed critically to the drafts and gave final approval for publication.

## Acknowledgements

We thank Wong Ching Ki and Tin Ka Yuen for their assistance with data collection and coding. We are grateful to Jean Leung of the South Lantau Buffalo Society for providing the buffalo demographic data. We sincerely thank Giacomo D’Ammando, Rene Corner-Thomas, Blair Costelloe, George Hodgson, and Tania Perroux for their valuable feedback on earlier versions of this manuscript. This research was funded by the Lantau Conservation Fund (Hong Kong SAR Government-funded program) (Grant ref. RE-2021-01).

## Conflict of interest statement

The authors declare no conflicts of interest.

## Statement of inclusion

The study brings together authors from several countries. The authors resided in the city in which the data were collected. The summary findings of the research will be made public and actively distributed (through project website, local social media groups, and non-government organisations) among residents of South Lantau in both English and Chinese languages.

## Data availability statement

Data and code will be made publicly available upon publication.

---

**Figure S1.**
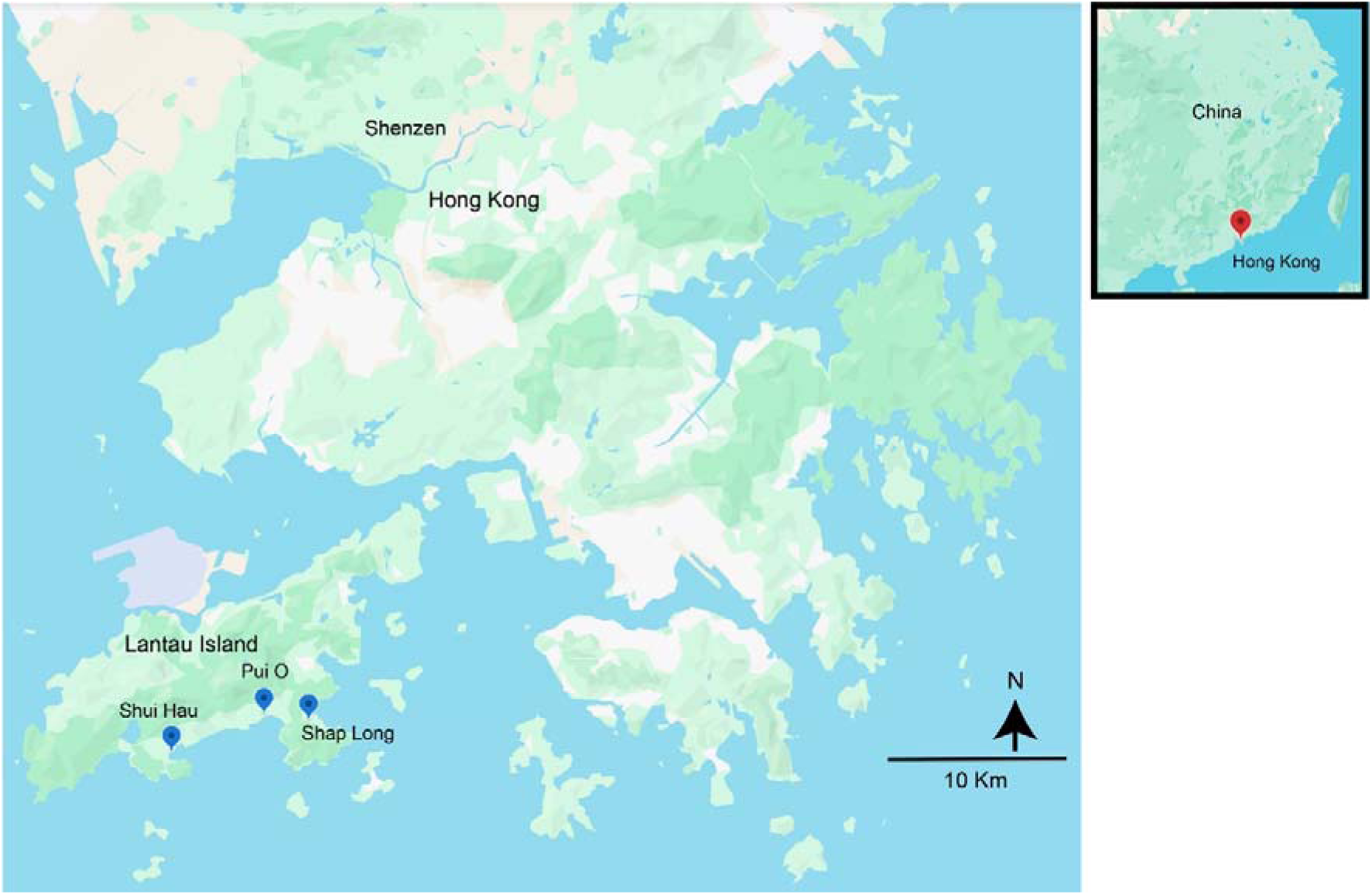
Geographic locations of the study area. A map of Hong Kong showing the geographic locations of the study area on Lantau Island: Pui O (Lo Wai Tsuen and Lo Uk Tsuen), Shap Long Kau Tsuen, and Shui Hau.

**Figure S2.**
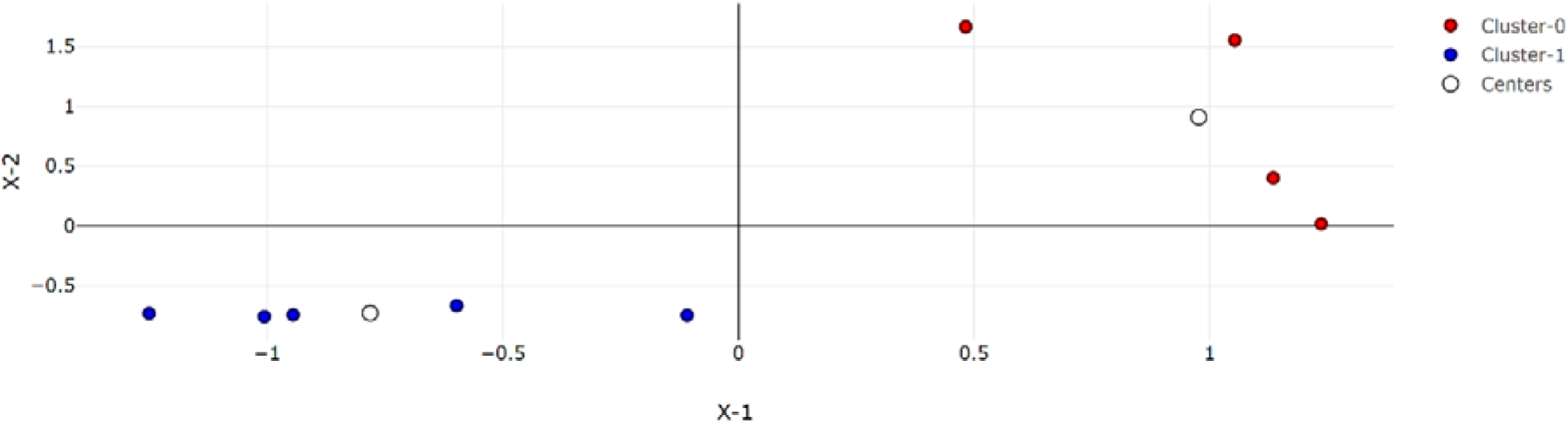
Results of K-means cluster analysis and the two distinct seasons based on publicly available weather data. Cluster 0 represents the months of July to October 2023, and Cluster 1 represents the months of November 2023 to March 2024. Cluster 0, or the wet season, had a median temperature of 29.4°C (range: 26.4°C – 30.1°C) and median rainfall of 391.2 mm (range: 175.2 mm – 546 mm). Cluster 1, or the dry season, had a median temperature of 19.4°C (range: 17.9°C – 23.5°C) and a median rainfall of 4.1 mm (range: 0.9 mm – 21.6 mm). k = 2 explained 80.19% of the variance of the data.

**Figure S3.**
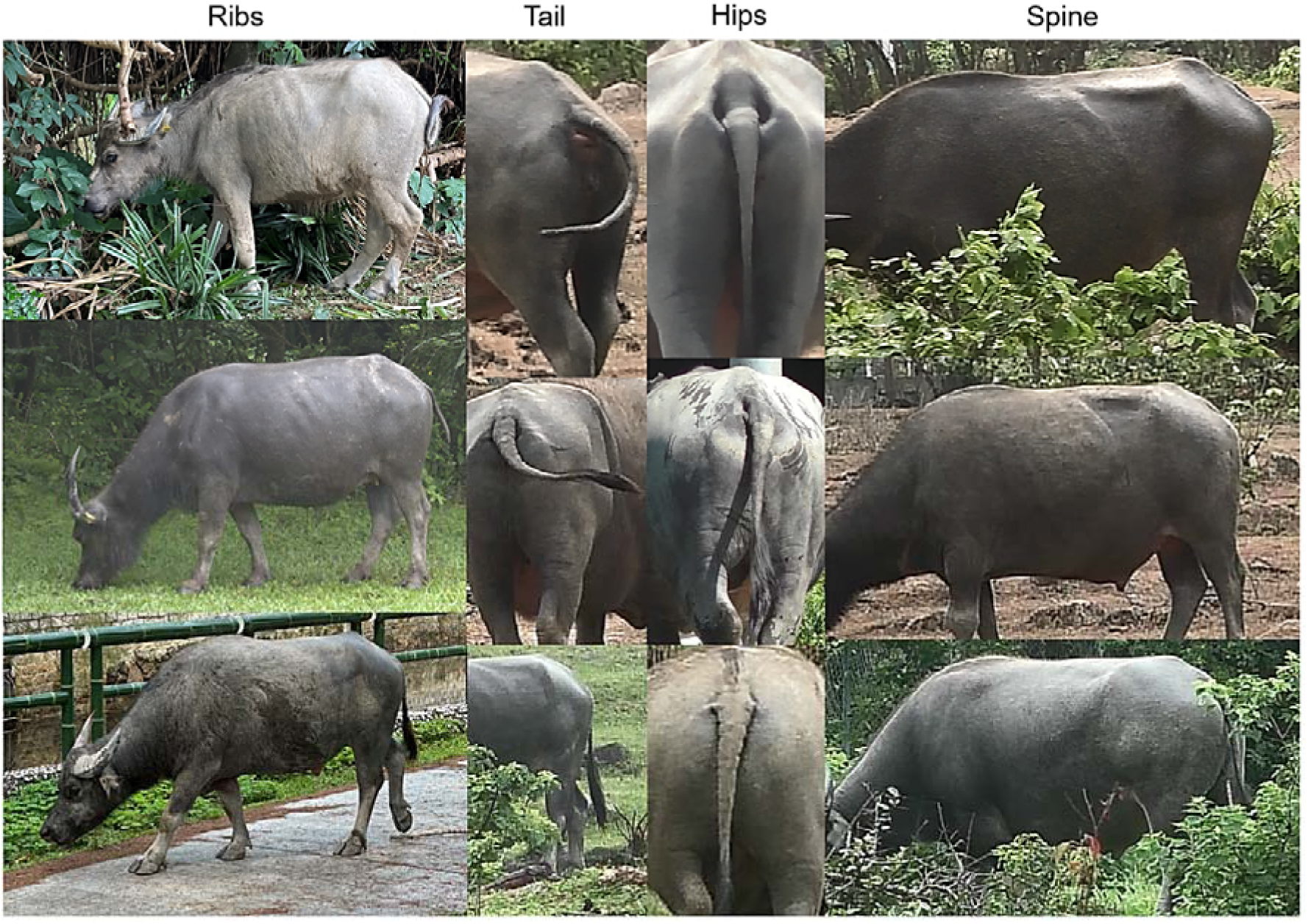
Different regions used for assessing buffalo body condition. The four different regions of the body are shown – ribs, tail, hips, and spine. For each region, conditions improve or scores increase from upper to lower photos.

**Figure S4:**
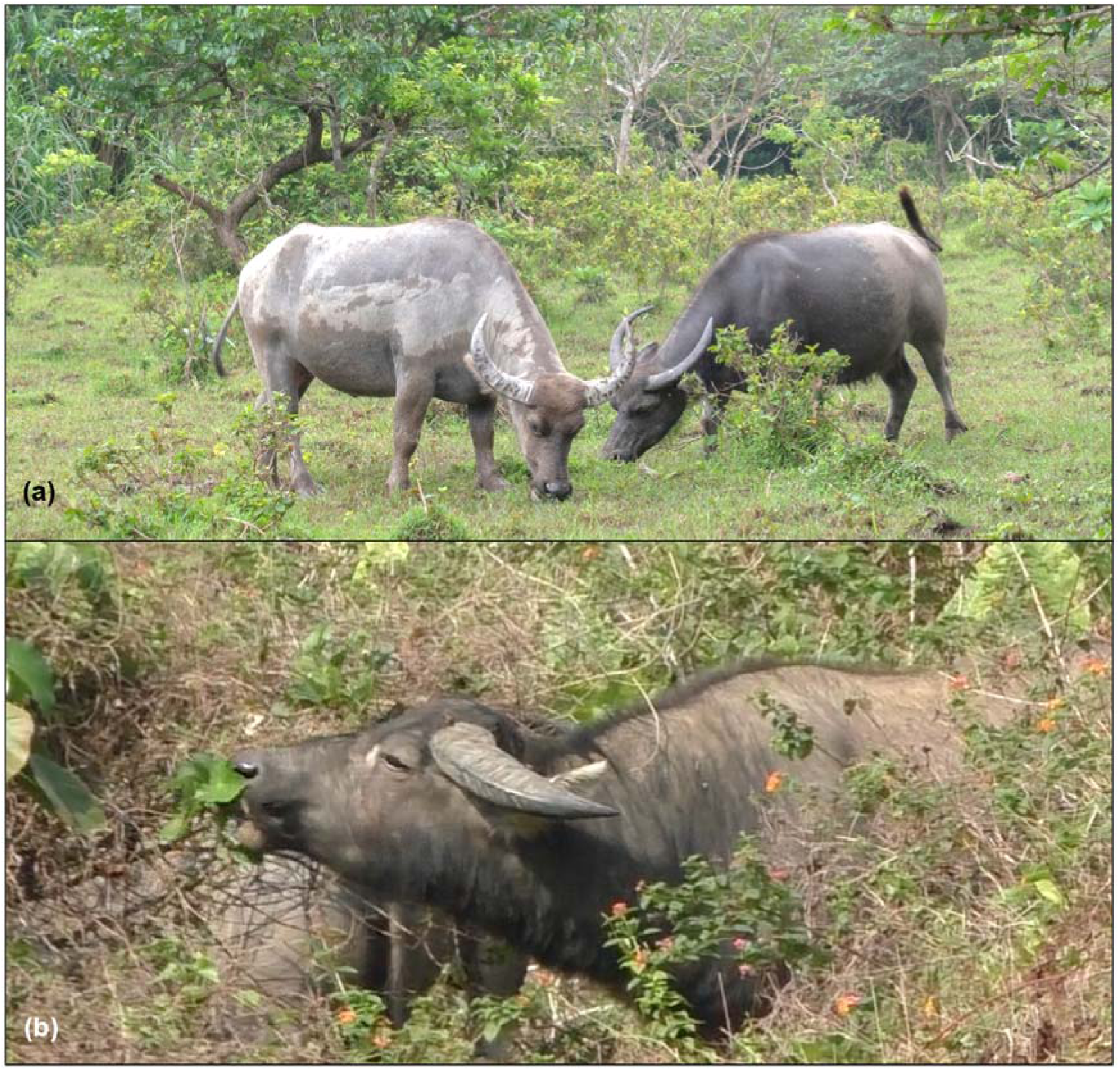
Feeding behaviour in buffalo. **(a)** Two female buffalo grazing on grass in Shui Hau, Lantau Island, Hong Kong and **(b)** A male buffalo browsing on leaves from shrubs in Shap Long Kau Tsuen, Lantau Island, Hong Kong [Photo credit: Debottam Bhattacharjee].

**Figure S5.**
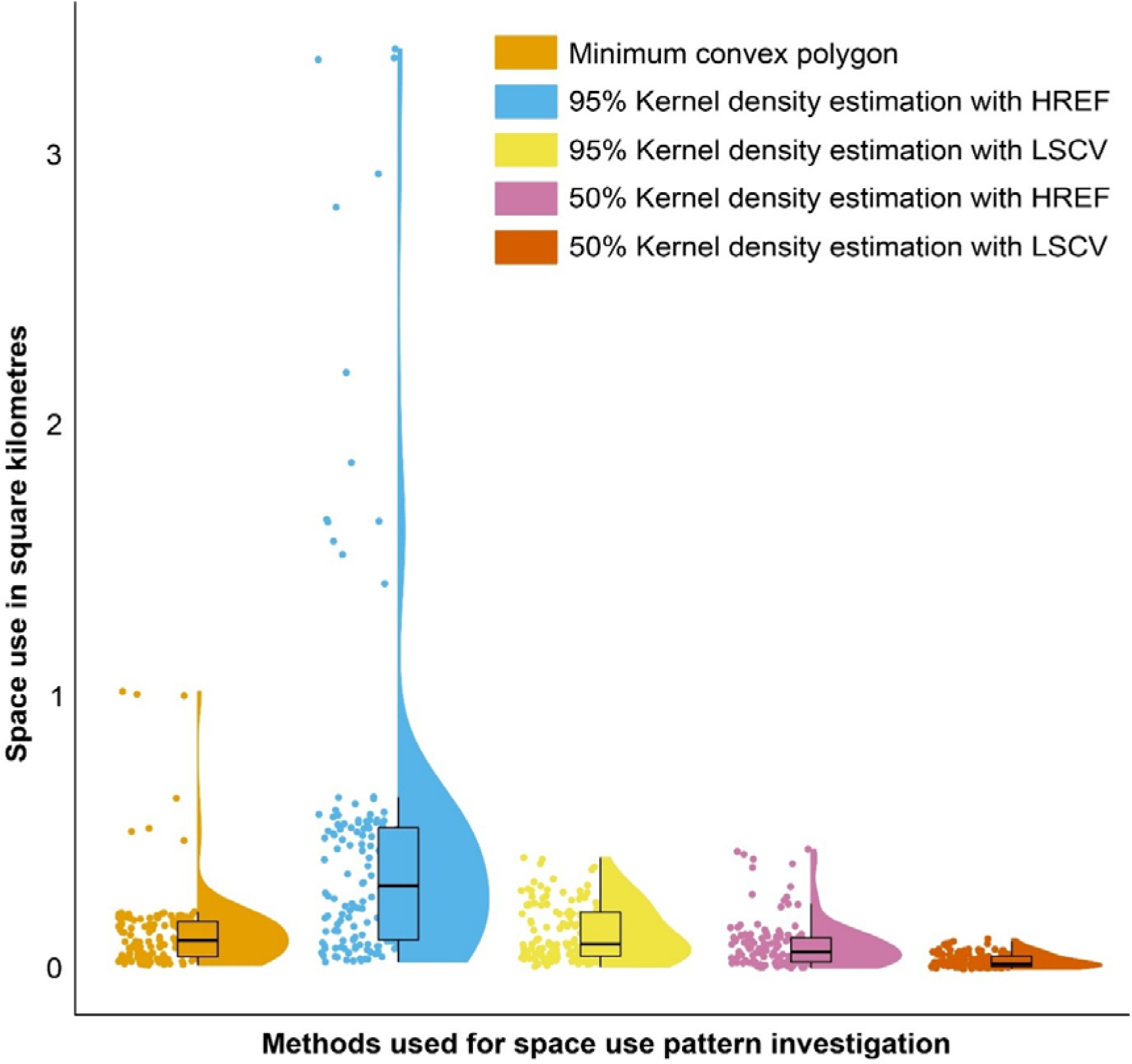
Buffalo space use pattern estimates based on different methods. Half-violin plots indicate the distribution of the data. Solid dots indicate the estimated area use values (based on 80 data points from wet and dry seasons) of the individuals (N = 71) calculated in square kilometres. The boxes illustrate the interquartile range, horizontal bars inside the boxes indicate median values, and whiskers indicate the range of the data.

**Figure S6.**
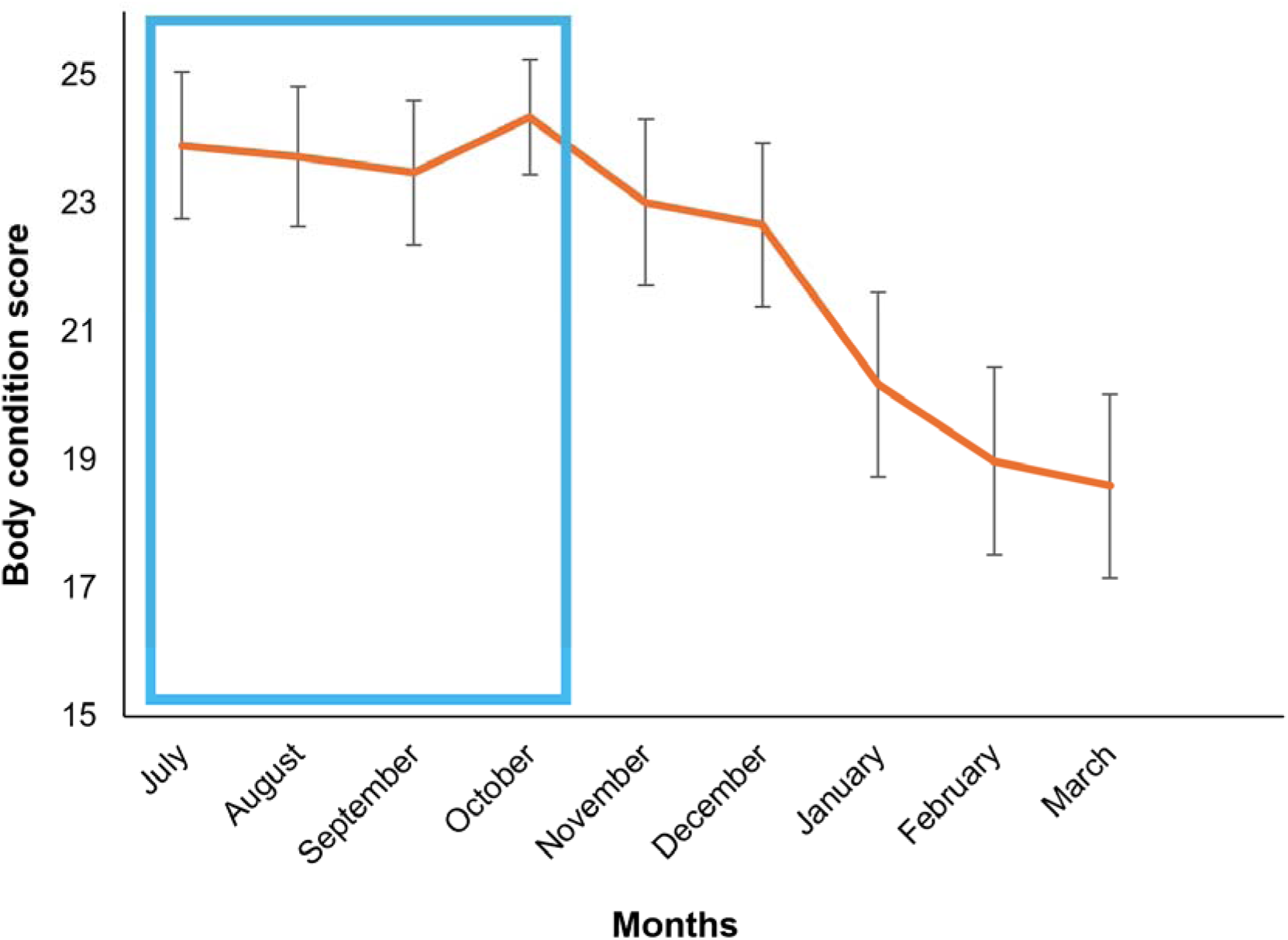
Monthly average body condition scores of buffalo. Mean (± standard deviation) body condition scores of buffalo (N = 71) from July 2023 to March 2024. Body condition scores were highest in the wet (July - October 2023) season (23.87 ± 0.88) and then declined during the dry (November 2023 – March 2024) season (19.20 ± 1.30). The wet season is highlighted by the blue box.

**Figure S7:**
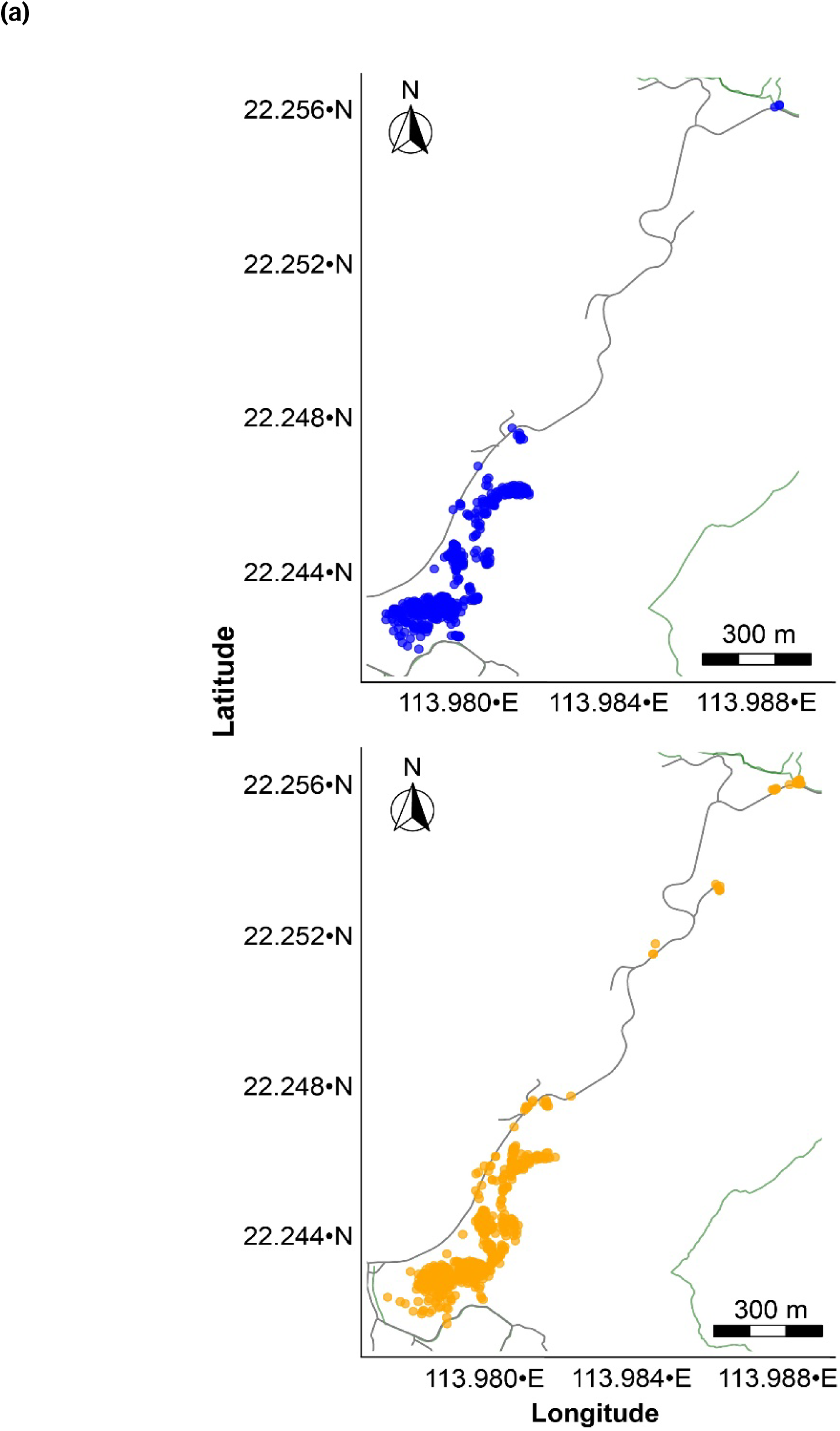
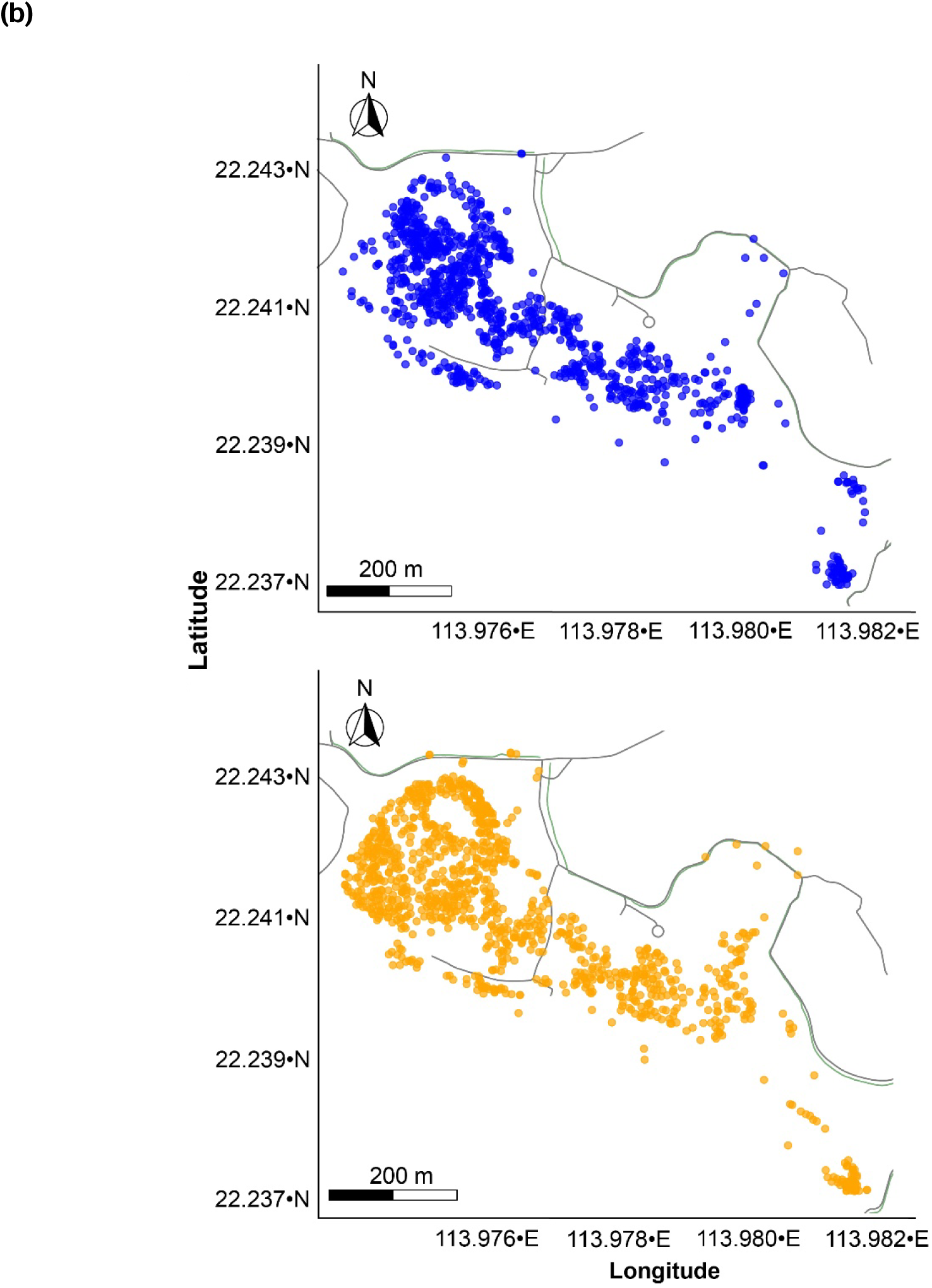
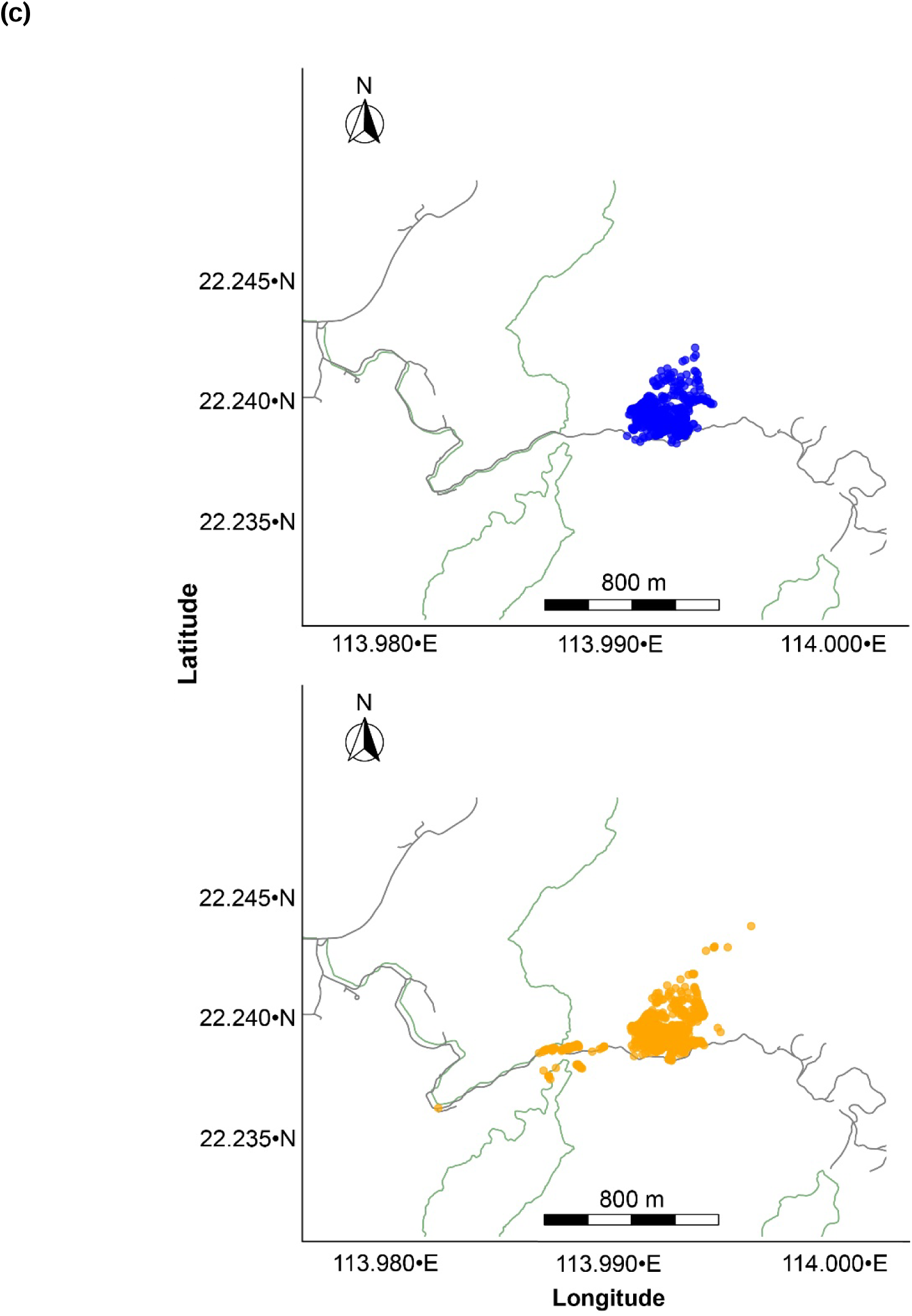
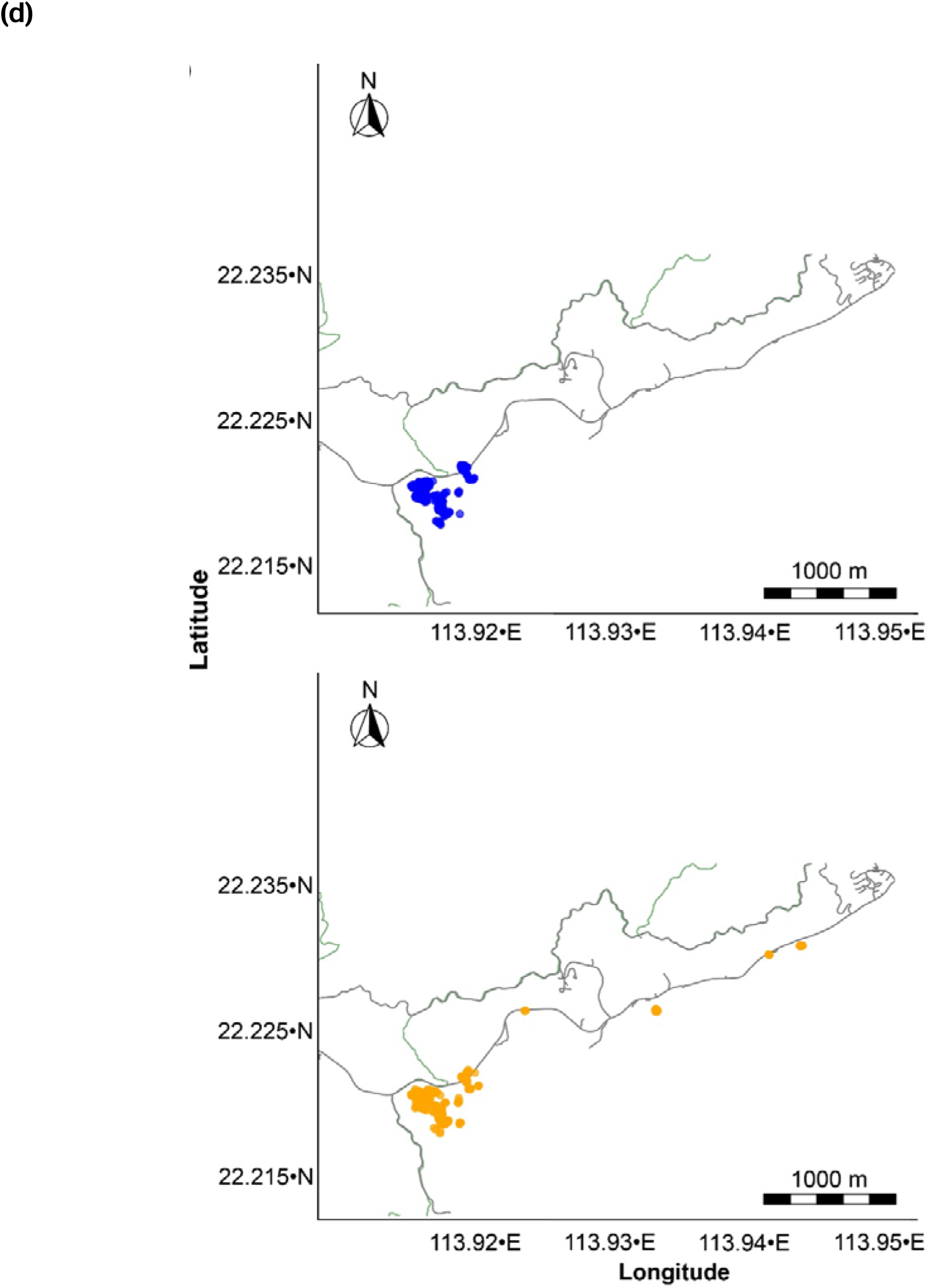
Location-specific distribution of buffalo during wet and dry seasons. The distribution of buffalo during wet and dry seasons in (a) Lo Wai Tsuen, (b) Lo Uk Tsuen, (c) Shap Long Kau Tsuen, and (d) Shui Hau. Blue dots show buffalo distribution during the wet season and orange dots show buffalo distribution during the dry season. Lantau Country Park borders, and major road networks are represented by green and black lines, respectively.

**Table S1.**
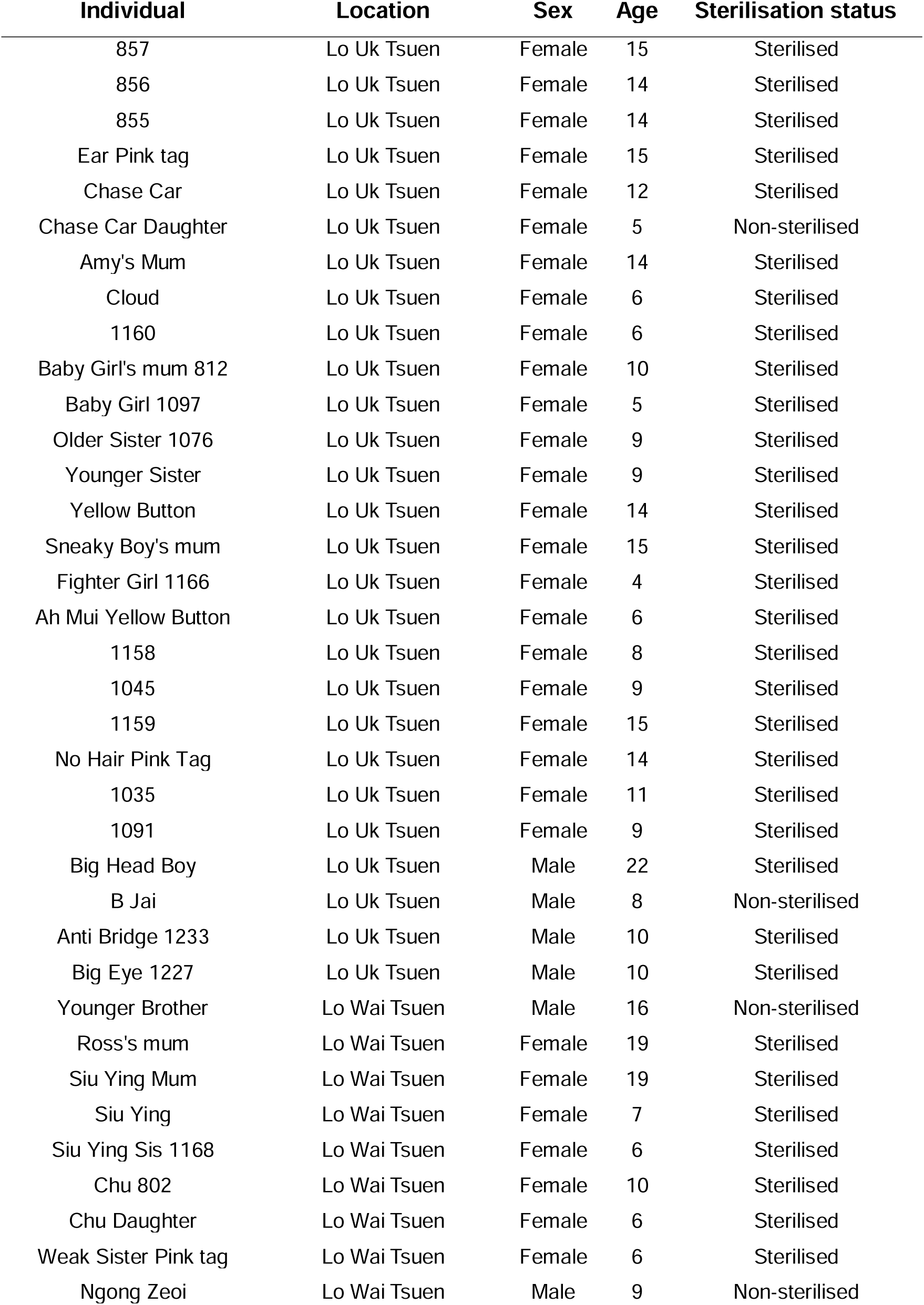
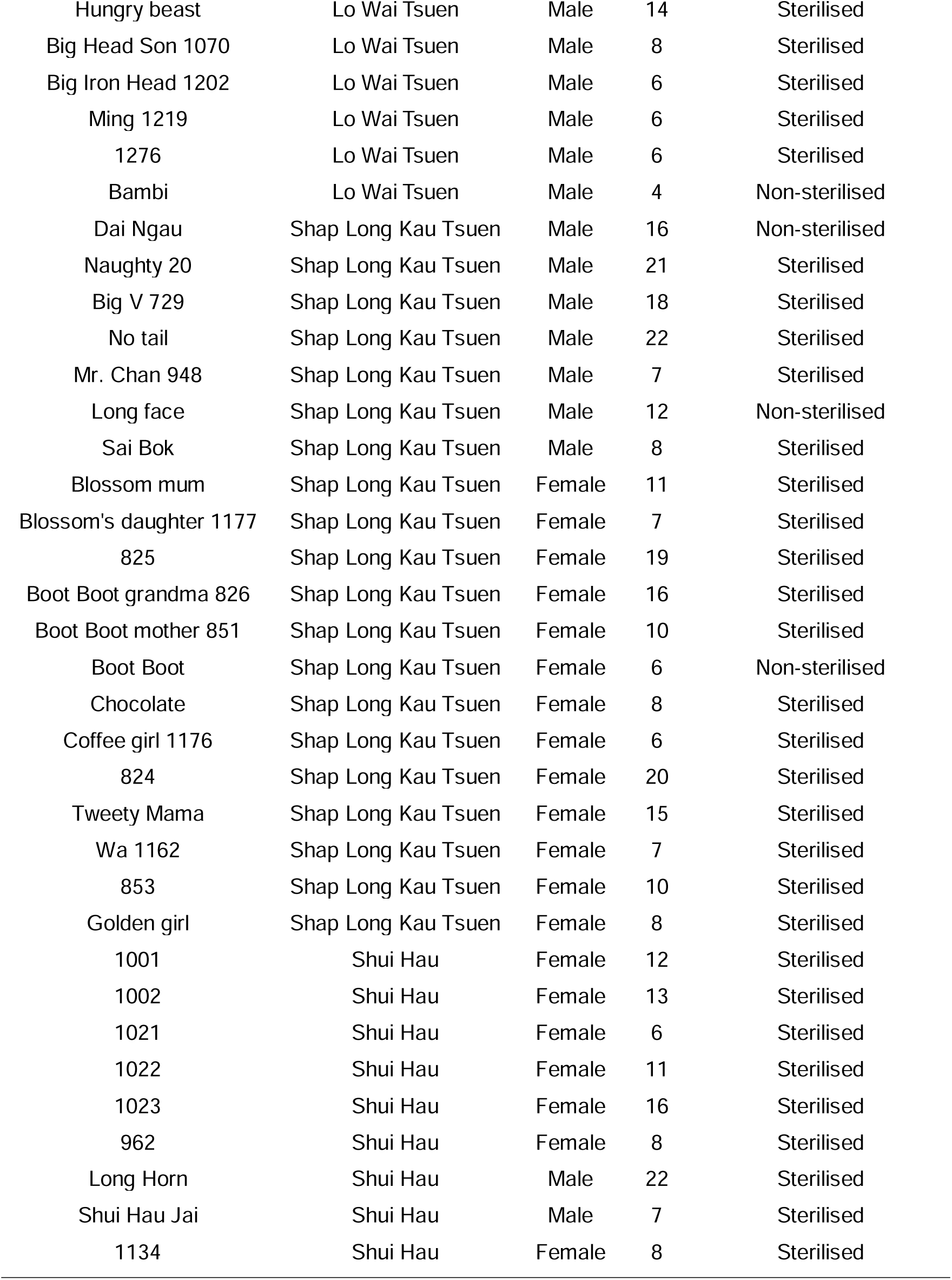
Summary of individual buffalo identity, location, sex, age, and sterilisation status.

**Table S2.** Summary of individual buffalo identity, location, and the percentage of visibility during first transects.

| Individual | Location | Percentage of visibility during first transects |
| --- | --- | --- |
| 857 | Lo Uk Tsuen | 100% |
| 856 | Lo Uk Tsuen | 100% |
| 855 | Lo Uk Tsuen | 100% |
| Ear Pink tag | Lo Uk Tsuen | 96.25% |
| Chase Car | Lo Uk Tsuen | 100% |
| Chase Car Daughter | Lo Uk Tsuen | 100% |
| Amy's Mum | Lo Uk Tsuen | 100% |
| Cloud | Lo Uk Tsuen | 100% |
| 1160 | Lo Uk Tsuen | 100% |
| Baby Girl's mum 812 | Lo Uk Tsuen | 100% |
| Baby Girl 1097 | Lo Uk Tsuen | 100% |
| Older Sister 1076 | Lo Uk Tsuen | 100% |
| Younger Sister | Lo Uk Tsuen | 100% |
| Yellow Button | Lo Uk Tsuen | 100% |
| Sneaky Boy's mum | Lo Uk Tsuen | 100% |
| Fighter Girl 1166 | Lo Uk Tsuen | 100% |
| Ah Mui Yellow Button | Lo Uk Tsuen | 100% |
| 1158 | Lo Uk Tsuen | 100% |
| 1045 | Lo Uk Tsuen | 100% |
| 1159 | Lo Uk Tsuen | 100% |
| No Hair Pink Tag | Lo Uk Tsuen | 98.75% |
| 1035 | Lo Uk Tsuen | 100% |
| 1091 | Lo Uk Tsuen | 100% |
| Big Head Boy | Lo Uk Tsuen | 100% |
| B Jai | Lo Uk Tsuen | 96.25% |
| Anti Bridge 1233 | Lo Uk Tsuen | 100% |
| Big Eye 1227 | Lo Uk Tsuen | 100% |
| Younger Brother | Lo Wai Tsuen | 98.75% |
| Ross's mum | Lo Wai Tsuen | 100% |
| Siu Ying Mum | Lo Wai Tsuen | 100% |
| Siu Ying | Lo Wai Tsuen | 100% |
| Siu Ying Sis 1168 | Lo Wai Tsuen | 100% |
| Chu 802 | Lo Wai Tsuen | 100% |
| Chu Daughter | Lo Wai Tsuen | 100% |
| Weak Sister Pink tag | Lo Wai Tsuen | 96.25% |
| Ngong Zeoi | Lo Wai Tsuen | 95% |
| Hungry beast | Lo Wai Tsuen | 95% |
| Big Head Son 1070 | Lo Wai Tsuen | 100% |
| Big Iron Head 1202 | Lo Wai Tsuen | 100% |
| Ming 1219 | Lo Wai Tsuen | 98.75% |
| 1276 | Lo Wai Tsuen | 100% |
| Bambi | Lo Wai Tsuen | 98.75% |
| Dai Ngau | Shap Long Kau Tsuen | 91.25% |
| Naughty 20 | Shap Long Kau Tsuen | 100% |
| Big V 729 | Shap Long Kau Tsuen | 100% |
| No tail | Shap Long Kau Tsuen | 100% |
| Mr. Chan 948 | Shap Long Kau Tsuen | 98.75% |
| Long face | Shap Long Kau Tsuen | 100% |
| Sai Bok | Shap Long Kau Tsuen | 87.5% |
| Blossom mum | Shap Long Kau Tsuen | 100% |
| Blossom's daughter 1177 | Shap Long Kau Tsuen | 100% |
| 825 | Shap Long Kau Tsuen | 100% |
| Boot Boot grandma 826 | Shap Long Kau Tsuen | 100% |
| Boot Boot mother 851 | Shap Long Kau Tsuen | 100% |
| Boot Boot | Shap Long Kau Tsuen | 100% |
| Chocolate | Shap Long Kau Tsuen | 100% |
| Coffee girl 1176 | Shap Long Kau Tsuen | 100% |
| 824 | Shap Long Kau Tsuen | 100% |
| Tweety Mama | Shap Long Kau Tsuen | 100% |
| Wa 1162 | Shap Long Kau Tsuen | 100% |
| 853 | Shap Long Kau Tsuen | 100% |
| Golden girl | Shap Long Kau Tsuen | 100% |
| 1001 | Shui Hau | 96.25% |
| 1002 | Shui Hau | 96.25% |
| 1021 | Shui Hau | 96.25% |
| 1022 | Shui Hau | 91.25% |
| 1023 | Shui Hau | 96.25% |
| 962 | Shui Hau | 96.25% |
| Long Horn | Shui Hau | 100% |
| Shui Hau Jai | Shui Hau | 98.75% |
| 1134 | Shui Hau | 95% |

## Notes

### Competing Interest Statement

The authors have declared no competing interest.

### Summary of Updates

Methods section updated to elaborate on observational processes, particularly body condition scoring section. Supplementary files updated.

## References

1. Abraham, J. O., Hempson, G. P., Faith, J. T., & Staver, A. C. (2022). Seasonal strategies differ between tropical and extratropical herbivores. Journal of Animal Ecology, 91(3), 681–692. 10.1111/1365-2656.13651

2. Abraham, J. O., Hempson, G. P., & Staver, A. C. (2019). Drought-response strategies of savanna herbivores. Ecology and Evolution, 9(12), 7047–7056. 10.1002/ece3.5270

3. AFCD. (2013). Stray cattle and buffalo management plan. Agriculture, Fisheries, and Conservation Department of Hong Kong. https://www.afcd.gov.hk/english/quarantine/cattlebuffalo.html

4. Aikens, E. O., Mysterud, A., Merkle, J. A., Cagnacci, F., Rivrud, I. M., Hebblewhite, M., Hurley, M. A., Peters, W., Bergen, S., De Groeve, J., Dwinnell, S. P. H., Gehr, B., Heurich, M., Hewison, A. J. M., Jarnemo, A., Kjellander, P., Kröschel, M., Licoppe, A., Linnell, J. D. C., … Kauffman, M. J. (2020). Wave-like patterns of plant phenology determine ungulate movement tactics. Current Biology, 30(17), 3444–3449.e4. 10.1016/j.cub.2020.06.032

5. Alapati, A., Kapa, S. R., Jeepalyam, S., Rangappa, S. M. P., & Yemireddy, K. R. (2010). Development of the body condition score system in Murrah buffaloes: validation through ultrasonic assessment of body fat reserves. Journal of Veterinary Science, 11(1), 1. 10.4142/jvs.2010.11.1.1

6. ASAB Ethical Committee, & ABS Animal Care Committee. (2022). Guidelines for the treatment of animals in behavioural research and teaching. Animal Behaviour, 183, I–XI. 10.1016/S0003-3472(21)00389-4

7. Beekman, J. H., & Prins, H. H. T. (1989). Feeding strategies of sedentary large herbivores in East Africa, with emphasis on the African buffalo, Syncerus coffer. African Journal of Ecology, 27(2), 129–147. 10.1111/j.1365-2028.1989.tb00937.x

8. Bergman, C. M., Fryxell, J. M., Gates, C. C., & Fortin, D. (2001). Ungulate foraging strategies: energy maximizing or time minimizing? Journal of Animal Ecology, 70(2), 289–300. 10.1111/j.1365-2656.2001.00496.x

9. Bhattacharjee, D., Flay, K. J., & McElligott, A. G. (2024). Personality homophily drives female friendships in a feral ungulate. IScience, 27(12), 111419. 10.1016/j.isci.2024.111419

10. Bonnot, N. C., Goulard, M., Hewison, A. J. M., Cargnelutti, B., Lourtet, B., Chaval, Y., & Morellet, N. (2018). Boldness-mediated habitat use tactics and reproductive success in a wild large herbivore. Animal Behaviour, 145, 107–115. 10.1016/j.anbehav.2018.09.013

11. Bora, J.K., Vardhan, V., Vijh, R.K., Deshmukh, A.V., Srinivas, Y., Mungi, N.A., Goswami, S., Jhala, H., Chauhan, J.S., Kumar, U., & Jhala, Y. (2024). Evaluating the potential for reintroducing the endangered wild water buffalo (Bubalus arnee) in Kanha National Park, central India. Restoration Ecology, 32(3), p.e14079.

12. Börger, L., Franconi, N., De Michele, G., Gantz, A., Meschi, F., Manica, A., Lovari, S., & Coulson, T. (2006). Effects of sampling regime on the mean and variance of home range size estimates. Journal of Animal Ecology, 75(6), 1393–1405. 10.1111/j.1365-2656.2006.01164.x

13. Bowman, D. M. J. S., Murphy, B. P., & McMahon, C. R. (2010). Using carbon isotope analysis of the diet of two introduced Australian megaherbivores to understand Pleistocene megafaunal extinctions. Journal of Biogeography, 37(3), 499–505. 10.1111/j.1365-2699.2009.02206.x

14. Bukombe, J., Kittle, A., Senzota, R., Mduma, S., Fryxell, J., & Sinclair, A. R. E. (2018). Resource selection, utilization and seasons influence spatial distribution of ungulates in the western Serengeti National Park. African Journal of Ecology, 56(1), 3–11. 10.1111/aje.12410

15. Bürkner, P.-C. (2018). Advanced Bayesian multilevel modeling with the R package brms. The R Journal, 10(1), 395. 10.32614/RJ-2018-017

16. Calenge, C. (2006). The package “adehabitat” for the R software: A tool for the analysis of space and habitat use by animals. Ecological Modelling, 197(3–4), 516–519. 10.1016/j.ecolmodel.2006.03.017

17. Campbell, H. A., Loewensteiner, D. A., Murphy, B. P., Pittard, S., & McMahon, C. R. (2021). Seasonal movements and site utilisation by Asian water buffalo (Bubalus bubalis) in tropical savannas and floodplains of northern Australia. Wildlife Research, 48(3), 230. 10.1071/WR20070

18. Carpio, A. J., Apollonio, M., & Acevedo, P. (2021). Wild ungulate overabundance in Europe: contexts, causes, monitoring and management recommendations. Mammal Review, 51(1), 95–108. 10.1111/mam.12221

19. Codron, D., Codron, J., Lee-Thorp, J. A., Sponheimer, M., De Ruiter, D., Sealy, J., Grant, R., & Fourie, N. (2007). Diets of savanna ungulates from stable carbon isotope composition of faeces. Journal of Zoology, 273(1), 21–29. 10.1111/j.1469-7998.2007.00292.x

20. Dagtekin, D., Ertürk, A., Sommer, S., Ozgul, A., & Soyumert, A. (2024). Seasonal habitat-use patterns of large mammals in a human-dominated landscape. Journal of Mammalogy, 105(1), 122–133. 10.1093/jmammal/gyad107

21. David Walter, W., Beringer, J., Hansen, L. P., Fischer, J. W., Millspaugh, J. J., & Vercauteren, K. C. (2011). Factors affecting space use overlap by white-tailed deer in an urban landscape. International Journal of Geographical Information Science, 25(3), 379–392. 10.1080/13658816.2010.524163

22. Depaoli, S., & van de Schoot, R. (2017). Improving transparency and replication in Bayesian statistics: The WAMBS-Checklist. Psychological Methods, 22(2), 240–261. 10.1037/met0000065

23. Dudgeon, D., & Corlett, R. T. (2011). The ecology and biodiversity of Hong Kong (Revised ed.). Cosmos Books & Lions Nature Education Foundation. https://doi.org/http://hdl.handle.net/10722/147078

24. Ezenwa, V. O., Jolles, A. E., & O’Brien, M. P. (2009). A reliable body condition scoring technique for estimating condition in African buffalo. African Journal of Ecology, 47(4), 476–481. 10.1111/j.1365-2028.2008.00960.x

25. Forrest, J., & Miller-Rushing, A. J. (2010). Toward a synthetic understanding of the role of phenology in ecology and evolution. Philosophical Transactions of the Royal Society B: Biological Sciences, 365(1555), 3101–3112. 10.1098/rstb.2010.0145

26. Gaidet, N., & Gaillard, J.-M. (2008). Density-dependent body condition and recruitment in a tropical ungulate. Canadian Journal of Zoology, 86(1), 24–32. 10.1139/Z07-111

27. Gaynor, K. M., Abrahms, B., Manlove, K. R., Oestreich, W. K., & Smith, J. A. (2024). Anthropogenic impacts at the interface of animal spatial and social behaviour. Philosophical Transactions of the Royal Society B: Biological Sciences, 379(1912). 10.1098/rstb.2022.0527

28. Hetem, R. S., Fuller, A., Maloney, S. K., & Mitchell, D. (2014). Responses of large mammals to climate change. Temperature, 1(2), 115–127. 10.4161/temp.29651

29. Holdo, R. M., Holt, R. D., & Fryxell, J. M. (2009). Opposing rainfall and plant nutritional gradients best explain the wildebeest migration in the Serengeti. The American Naturalist, 173(4), 431–445. 10.1086/597229

30. Honda, T., Iijima, H., Tsuboi, J., & Uchida, K. (2018). A review of urban wildlife management from the animal personality perspective: The case of urban deer. Science of The Total Environment, 644, 576–582. 10.1016/j.scitotenv.2018.06.335

31. Hong Kong Observatory. (2024). Monthly weather summary. https://www.hko.gov.hk/en/wxinfo/pastwx/mws/mws.htm

32. Kartzinel, T. R., Chen, P. A., Coverdale, T. C., Erickson, D. L., Kress, W. J., Kuzmina, M. L., Rubenstein, D. I., Wang, W., & Pringle, R. M. (2015). DNA metabarcoding illuminates dietary niche partitioning by African large herbivores. Proceedings of the National Academy of Sciences of the United States of America, 112(26), 8019–8024. 10.1073/pnas.1503283112

33. Kaszta, Ż., Marino, J., Ramoelo, A., & Wolff, E. (2016). Bulk feeder or selective grazer: African buffalo space use patterns based on fine-scale remotely sensed data on forage quality and quantity. Ecological Modelling, 323, 115–122. 10.1016/j.ecolmodel.2015.12.006

34. Koshida, C., & Katayama, N. (2018). Meta-analysis of the effects of rice-field abandonment on biodiversity in Japan. Conservation Biology, 32(6), 1392–1402. 10.1111/cobi.13156

35. Laver, P. N., & Kelly, M. J. (2008). A critical review of home range studies. The Journal of Wildlife Management, 72(1), 290–298. 10.2193/2005-589

36. Lemoine, N. P. (2019). Moving beyond noninformative priors: why and how to choose weakly informative priors in Bayesian analyses. Oikos, 128(7), 912–928. 10.1111/oik.05985

37. Lewis, S. L., & Maslin, M. A. (2015). Defining the Anthropocene. Nature, 519(7542), 171–180. 10.1038/nature14258

38. Lichti, N. I., & Swihart, R. K. (2011). Estimating utilization distributions with kernel versus local convex hull methods. The Journal of Wildlife Management, 75(2), 413–422. 10.1002/jwmg.48

39. Lundgren, E. J., Bergman, J., Trepel, J., le Roux, E., Monsarrat, S., Kristensen, J. A., Pedersen, R. Ø., Pereyra, P., Tietje, M., & Svenning, J.-C. (2024). Functional traits—not nativeness—shape the effects of large mammalian herbivores on plant communities. Science, 383(6682), 531–537. 10.1126/science.adh2616

40. Makowski, D., Ben-Shachar, M., Patil, I., & Lüdecke, D. (2020). Methods and algorithms for correlation analysis in R. Journal of Open Source Software, 5(51), 2306. 10.21105/joss.02306

41. McCain, C. M., & King, S. R. B. (2014). Body size and activity times mediate mammalian responses to climate change. Global Change Biology, 20(6), 1760–1769. 10.1111/gcb.12499

42. McElreath, R. (2020). *Statistical Rethinking* (2nd Edition). Chapman and Hall/CRC. 10.1201/9780429029608

43. Mihailou, H., & Massaro, M. (2021). An overview of the impacts of feral cattle, water buffalo and pigs on the savannas, wetlands and biota of northern Australia. Austral Ecology, 46(5), 699–712. 10.1111/aec.13046

44. Minervino, A. H. H., Zava, M., Vecchio, D., & Borghese, A. (2020). Bubalus bubalis: A Short Story. Frontiers in Veterinary Science, 7. 10.3389/fvets.2020.570413

45. Monestier, C., Gilot-Fromont, E., Morellet, N., Debeffe, L., Cebe, N., Merlet, J., Picot, D., Rames, J.-L., Hewison, A. J. M., & Verheyden, H. (2016). Individual variation in an acute stress response reflects divergent coping strategies in a large herbivore. Behavioural Processes, 132, 22–28. 10.1016/j.beproc.2016.09.004

46. Monsarrat, S., Jarvie, S., & Svenning, J.-C. (2019). Anthropocene refugia: integrating history and predictive modelling to assess the space available for biodiversity in a human-dominated world. Philosophical Transactions of the Royal Society B: Biological Sciences, 374(1788), 20190219. 10.1098/rstb.2019.0219

47. Natuhara, Y. (2013). Ecosystem services by paddy fields as substitutes of natural wetlands in Japan. Ecological Engineering, 56, 97–106. 10.1016/j.ecoleng.2012.04.026

48. Niu, S. Q., & Dudgeon, D. (2011). Environmental flow allocations in monsoonal Hong Kong. Freshwater Biology, 56(6), 1209–1230. 10.1111/j.1365-2427.2010.02558.x

49. O’Donnell, K., & delBarco-Trillo, J. (2020). Changes in the home range sizes of terrestrial vertebrates in response to urban disturbance: a meta-analysis. Journal of Urban Ecology, 6(1). 10.1093/jue/juaa014

50. Ogutu, J. O., Piepho, H.-P., & Dublin, H. T. (2014). Reproductive seasonality in African ungulates in relation to rainfall. Wildlife Research, 41(4), 323. 10.1071/WR13211

51. Owen-Smith, N., Mason, D. R., & Ogutu, J. O. (2005). Correlates of survival rates for 10 African ungulate populations: density, rainfall and predation. Journal of Animal Ecology, 74(4), 774–788. 10.1111/j.1365-2656.2005.00974.x

52. Owen-Smith, N., & Ogutu, J. O. (2013). Controls over reproductive phenology among ungulates: allometry and tropical-temperate contrasts. Ecography, 36(3), 256–263. 10.1111/j.1600-0587.2012.00156.x

53. Parker, K. L., Barboza, P. S., & Gillingham, M. P. (2009). Nutrition integrates environmental responses of ungulates. Functional Ecology, 23(1), 57–69. 10.1111/j.1365-2435.2009.01528.x

54. Pascual-Rico, R., Morales-Reyes, Z., Aguilera-Alcalá, N., Olszańska, A., Sebastián-González, E., Naidoo, R., Moleón, M., Lozano, J., Botella, F., von Wehrden, H., Martín-López, B., & Sánchez-Zapata, J. A. (2021). Usually hated, sometimes loved: A review of wild ungulates’ contributions to people. Science of The Total Environment, 801, 149652. 10.1016/j.scitotenv.2021.149652

55. Pebesma, E. (2018). Simple features for R: standardized support for spatial vector data. The R Journal, 10(1), 439. 10.32614/RJ-2018-009

56. Petty, A. M., Werner, P. A., Lehmann, C. E. R., Riley, J. E., Banfai, D. S., & Elliott, L. P. (2007). Savanna responses to feral buffalo in Kakadu national park, Australia. Ecological Monographs, 77(3), 441–463. 10.1890/06-1599.1

57. Pike, K. N., Perry, J., Vanderduys, E., Arnould, J. P. Y., & Hoskins, A. (2024). Love thy neighbour: Feral buffalos show greater space use, resource overlap and encounters during the wet season in the Northern Territory. Ecology and Evolution, 14(10). 10.1002/ece3.70345

58. Pokharel, S. S., Seshagiri, P. B., & Sukumar, R. (2017). Assessment of season-dependent body condition scores in relation to faecal glucocorticoid metabolites in free-ranging Asian elephants. Conservation Physiology, 5(1). 10.1093/conphys/cox039

59. Polfus, J. L., & Krausman, P. R. (2012). Impacts of residential development on ungulates in the Rocky Mountain West. Wildlife Society Bulletin, 36(4), 647–657. 10.1002/wsb.185

60. Powell, D. M., Beetem, D., Breitigan, R., Eyres, A., & Speeg, B. (2024). A perspective on ungulate management and welfare assessment across the traditional zoo to large landscape spectrum. Zoo Biology, 43(1), 5–14. 10.1002/zoo.21772

61. Powell, R. A. (2000). Animal home ranges and territories and home range estimators. In L. Boitani & T. K. Fuller (Eds.), Research Techniques in Animal Ecology: Controversies and Consequences (pp. 65–110). Columbia University. https://www.researchgate.net/publication/304436028

62. R Development Core Team. (2019). R Core Team (2020). R: A language and environment for statistical computing. R Foundation for Statistical Computing, Vienna, Austria. URL https://www.R-project.org/. In R Foundation for Statistical Computing (Vol. 2).

63. Ranglack, D. H., & du Toit, J. T. (2015). Wild bison as ecological indicators of the effectiveness of management practices to increase forage quality on open rangeland. Ecological Indicators, 56, 145–151. 10.1016/j.ecolind.2015.04.009

64. Rödel, H. G., Ibler, B., Ozogány, K., & Kerekes, V. (2023). Age-specific effects of density and weather on body condition and birth rates in a large herbivore, the Przewalski’s horse. Oecologia, 203(3–4), 435–451. 10.1007/s00442-023-05477-9

65. Rodríguez-Sánchez, F. (2024). DHARMa.helpers: Helper functions to check models not (yet) directly supported by DHARMa (R package version 0.0.2). https://github.com/Pakillo/DHARMa.helpers

66. Roger, S. B., Edzer, P., & Virgilio, G.-R. (2013). *Applied spatial data analysis with R* (Second). Springer. https://asdar-book.org/

67. Roug, A., Muse, E. A., Clifford, D. L., Larsen, R., Paul, G., Mathayo, D., Mpanduji, D., Mazet, J. A. K., Kazwala, R., Kiwango, H., & Smith, W. (2020). Seasonal movements and habitat use of African buffalo in Ruaha National Park, Tanzania. BMC Ecology, 20(1), 6. 10.1186/s12898-020-0274-4

68. Schirmer, A., Herde, A., Eccard, J. A., & Dammhahn, M. (2019). Individuals in space: personality-dependent space use, movement and microhabitat use facilitate individual spatial niche specialization. Oecologia, 189(3), 647–660. 10.1007/s00442-019-04365-5

69. Seaman, D. E., Millspaugh, J. J., Kernohan, B. J., Brundige, G. C., Raedeke, K. J., & Gitzen, R. A. (1999). Effects of sample size on kernel home range estimates. The Journal of Wildlife Management, 63(2), 739. 10.2307/3802664

70. So, K. Y. K., & Dudgeon, D. (2020). Conservation management of abandoned paddy fields in Asia: Semi-natural marshes with low-intensity bovid grazing have higher biodiversity. Aquatic Conservation: Marine and Freshwater Ecosystems, 30(10), 1934–1944. 10.1002/aqc.3442

71. Spiegel, O., Leu, S. T., Bull, C. M., & Sih, A. (2017). What’s your move? Movement as a link between personality and spatial dynamics in animal populations. Ecology Letters, 20(1), 3–18. 10.1111/ele.12708

72. Staver, A. C., & Hempson, G. P. (2020). Seasonal dietary changes increase the abundances of savanna herbivore species. Science Advances, 6(40). 10.1126/sciadv.abd2848

73. Stephenson, T. R., German, D. W., Cassirer, E. F., Walsh, D. P., Blum, M. E., Cox, M., Stewart, K. M., & Monteith, K. L. (2020). Linking population performance to nutritional condition in an alpine ungulate. Journal of Mammalogy, 101(5), 1244–1256. 10.1093/jmammal/gyaa091

74. Stiegler, J., Lins, A., Dammhahn, M., Kramer-Schadt, S., Ortmann, S., & Blaum, N. (2022). Personality drives activity and space use in a mammalian herbivore. Movement Ecology, 10(1), 33. 10.1186/s40462-022-00333-6

75. Tang, K.-L. (2017). Whose development? Transnational capitalism and the homogenisation of space. In Encountering Development in the Age of Global Capitalism (pp. 147–164). Springer Singapore. 10.1007/978-981-10-5120-3_6

76. Thackeray, S. J., Henrys, P. A., Hemming, D., Bell, J. R., Botham, M. S., Burthe, S., Helaouet, P., Johns, D. G., Jones, I. D., Leech, D. I., Mackay, E. B., Massimino, D., Atkinson, S., Bacon, P. J., Brereton, T. M., Carvalho, L., Clutton-Brock, T. H., Duck, C., Edwards, M., … Wanless, S. (2016). Phenological sensitivity to climate across taxa and trophic levels. Nature, 535(7611), 241–245. 10.1038/nature18608

77. Varpe, Ø. (2017). Life History Adaptations to Seasonality. Integrative and Comparative Biology, 57(5), 943–960. 10.1093/icb/icx123

78. Webb, W. C., Marzluff, J. M., & Hepinstall-Cymerman, J. (2011). Linking resource use with demography in a synanthropic population of common ravens. Biological Conservation, 144(9), 2264–2273. 10.1016/j.biocon.2011.06.001

79. Wevers, J., Fattebert, J., Casaer, J., Artois, T., & Beenaerts, N. (2020). Trading fear for food in the Anthropocene: How ungulates cope with human disturbance in a multi-use, suburban ecosystem. Science of The Total Environment, 741, 140369. 10.1016/j.scitotenv.2020.140369

80. Williams, C. M., Ragland, G. J., Betini, G., Buckley, L. B., Cheviron, Z. A., Donohue, K., Hereford, J., Humphries, M. M., Lisovski, S., Marshall, K. E., Schmidt, P. S., Sheldon, K. S., Varpe, Ø., & Visser, M. E. (2017). Understanding evolutionary impacts of seasonality: An introduction to the symposium. Integrative and Comparative Biology, 57(5), 921–933. 10.1093/icb/icx122

81. Williams, C. T., Chmura, H. E., Deal, C. K., & Wilsterman, K. (2022). Sex-differences in phenology: A Tinbergian perspective. Integrative and Comparative Biology, 62(4), 980– 997. 10.1093/icb/icac035

82. Wittemyer, G., Getz, W. M., Vollrath, F., & Douglas-Hamilton, I. (2007). Social dominance, seasonal movements, and spatial segregation in African elephants: a contribution to conservation behavior. Behavioral Ecology and Sociobiology, 61(12), 1919–1931. 10.1007/s00265-007-0432-0

83. Woodworth, B. K., Norris, D. R., Graham, B. A., Kahn, Z. A., & Mennill, D. J. (2018). Hot temperatures during the dry season reduce survival of a resident tropical bird. Proceedings of the Royal Society B: Biological Sciences, 285(1878), 20180176. 10.1098/rspb.2018.0176

84. WWF-Hong Kong. (2021). Preserving our natural wealth of Southern Lantau Island of Hong Kong. https://wwfhk.awsassets.panda.org/downloads/wwf_south_lantau_watershed_analysis_27sept2021.pdf

85. Yang, D., Bhattacharjee, D., Flay, K. J., Wang, Y., Mumby, H. S., & McElligott, A. G. (2024). Public attitudes and values regarding a semi-urban feral megaherbivore. SocArXiv. 10.31235/osf.io/yu9rj

86. Young, K. D., Ferreira, S. M., & Van Aarde, R. J. (2009). Elephant spatial use in wet and dry savannas of southern Africa. Journal of Zoology, 278(3), 189–205. 10.1111/j.1469-7998.2009.00568.x

